# Novel microbial syntrophies identified by longitudinal metagenomics

**DOI:** 10.1101/2021.07.05.451125

**Authors:** Sebastien Raguideau, Anna Trego, Fred Farrell, Gavin Collins, Christopher Quince, Orkun S Soyer

## Abstract

Identifying species interactions in a microbial community and how this relates to community function is a key challenge. Towards addressing this challenge, we present here an extensive genome-resolved, longitudinal dataset and associated metadata. We collected weekly samples of microbial communities and recorded operating conditions from industrial methane producing anaerobic digestion reactors for a year. This allowed us to recover 2240 dereplicated metagenome assembled genomes (dMAGs), together with their coverage dynamics and functional annotations from which functional traits were inferred. Of these dMAGs, 1910 were novel species, with 22 representing novel orders and classes. Methanogenic communities are expected to be strongly structured by syntrophic and other associations between the methanogens and syntrophs that produce their substrates. We identified 450 potential syntrophic dMAGs by searching for pairs of methanogenic and non-methanogenic dMAGs that had highly correlated time-series. Genomes of potential syntrophs were enriched for oxidoreductases and sugar transport genes and there was a strong taxonomic signal in their associations with methanogens. Of particular note, we found that Bathyarchaeiea associated specifically with methanogens from the Thermoplasmata, and Thermococci classes. Same syntrophic associations were only rarely observed across multiple reactors, suggesting that syntrophies might be facultative, with particular strains within a species forming syntrophic associations only sometimes and not necessarily always with the same methanogenic partner. The presented results show that longitudinal metagenomics is a highly valuable approach for identifying species and their interactions in microbial communities.

**One Sentence Summary:** Longitudinal study of microbial communities identifies novel species and predicts their interactions and role in community function.

## INTRODUCTION

Over the last decade, genome-resolved metagenome sequencing of samples from diverse environments generated thousands of metagenome-assembled genomes (MAGs) (*1*). These included new diversity in the Candidate Phlyla Radiation (CPR), bacteria, and archaea, including novel phyla (*2-6*), and reshaped our understanding of microbial phylogeny (*7*). Many novel species identified through metagenomics lack cultivated representatives, raising the question of their functional role and interactions within the communities that they reside in. Functional capacities of MAGs can be resolved to some extent through analysis of genomic content. Thus, several novel MAGs, including some CPRs, were shown to harbour genes associated with key processes including organic degradation (*2,8,9*) and nitrogen, and sulfur, cycling (*10-13*).

While genome content can suggest potential functional capacities in novel species, it cannot predict which species interactions are present in a community and how these might dynamically change to impact overall community function. One way to try and answer these questions is to combine functional analysis of MAGs with their abundances over time and with associated measurements of community outputs and the environmental parameters (*14*). Such combined longitudinal studies of microbial communities and metadata from the same system are still not that common, but provide useful insights whenever conducted. For example, using a small number of samples collected over a season, and combining these with chemical analyses, it was possible to identify key species and environmentally driven changes within animal rumen and industrial fermentation communities (*15-17*). It has been also possible to use temporal samples on their own, without any metadata, to predict possible interactions in a subsurface community (*18,19*), as well as in an infant gut (*20*).

Here, we report results from a comprehensive, longitudinal study of methanogenic microbial communities found in industrial anaerobic digestion (AD) bioreactors. Formation of methane from organic matter in AD reactors is a community-level process, since methanogenic substrates are the metabolic by-products of fermenters. Many fermentation pathways are prone to thermodynamic inhibition due to product accumulation (*21, 22*), making syntrophic associations between methanogens and fermenters essential for the completion of the AD process (*23-25*). This, in turn, allows for methane production to be used as a proxy for community stability and performance (*26*). These features, along with regular monitoring of methane production and other environmental parameters, make AD systems an ideal choice for a longitudinal analysis. We find that the presented high-resolution, longitudinal, genome-resolved metagenomics approach allows identification of 2240 species and 518 potential interactions in AD reactor communities and reveals associations between environmental parameters, methane production, and functional trait dynamics. These findings show that the design and conduct of longitudinal studies can provide a highly useful tool to understand and manage microbial communities.

## RESULTS AND DISCUSSION

We have sampled DNA and collected metadata from industrial AD reactors on a weekly basis for over 30 weeks and a maximum of 39 weeks (see *Methods* and Table *S1*). The reactor metadata was composed of temperature, pH, methane and hydrogen sulphide concentration, gas and energy production rate, and feed rate. All of the resulting sequence data from this project are deposited on the European Nucleotide Archive (see *Methods*).

### Sequence analysis identifies 2240 species, contributing to significant expansion of existing phyla and identification of new classes and orders

We performed reactor-wise co-assembly and binning of our sequence data, which on average, comprised 41 million reads per sample (see *Methods* and Fig. S1). This resulted in 5303 individual MAGs with at least 75% of their single-copy genes in single-copy. These MAGs were than clustered using a threshold of 99% similarity over at least 10% of their sequence, resulting in 2240 dereplicated MAGs (dMAGs). These dMAGs had a median completion of 93% and a median contamination of 1.4% (Fig. S2). We annotated each dMAG for genomic content by calling KEGG orthologs (KOs) (see *Methods* and Fig. S2 for a summary). Furthermore, each dMAG could be reliably assigned a normalised coverage that was proportional to their relative genome copy number in each sample (see *Methods* and Fig. S1). This was then used for time series abundance and correlation analyses.

To taxonomically classify dMAGs, we used the GTDB-Tk (*27*), that is based on over 190,000 known genomes available at the Genome Taxonomy Database (GTDB) (*28*). In addition, we have constructed a *de novo* phylogenetic tree of all dMAGs, using a maximum likelihood method (see *Methods*). When failing direct assignment to a known species GTDB-Tk provides a standardized relative evolutionary divergence (RED), an empirical quantification of taxonomic novelty that is shown to have a mean of 0.32, 0.48, and 0.63 at phylum, class, and order levels respectively (*29*). The taxonomic assignment and RED value of dMAGs found in this study – 2195 Bacteria and 45 Archaea –are summarised on Fig. 1 and 2.

**Figure 1.**
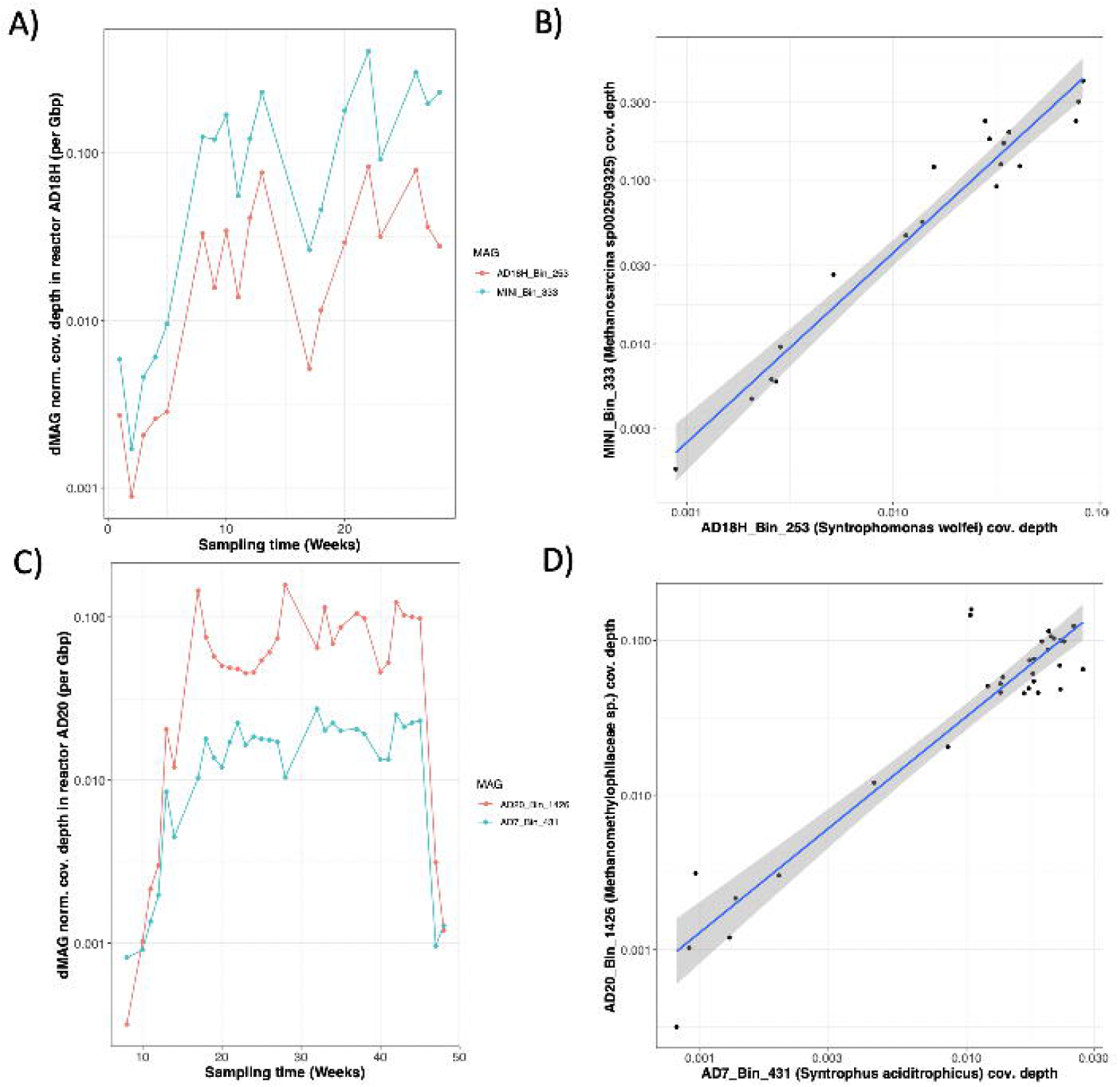
Phylogenetic relations and novelty of bacterial dMAGs identified in this study. **(A)** A phylogenetic tree created with all 2,195 bacterial dMAGs and representatives from GTDB, using a maximum likelihood approach (see *Methods*). Only the dMAGs are shown, along with their RED value encoded in grey shading on the outer ring (see legend). Branch colours indicate major phyla, as shown in the legend and with the “PVC” branch including Verrucomicrobiota, Planctomycetota, SZUA-79, Omnitrophota, Poribacteria, and UBP3. **(B)** The fold expansion of the representation in the GTDB database for those phyla, where the expansion was above 0.3. **(C)** The distribution of RED values for the 2,195 bacterial dMAGs. Stacked bars show the different phyla in each bin. For (B) and (C), the same colouring scheme of phyla is used, as in (A).

**Figure 2.**
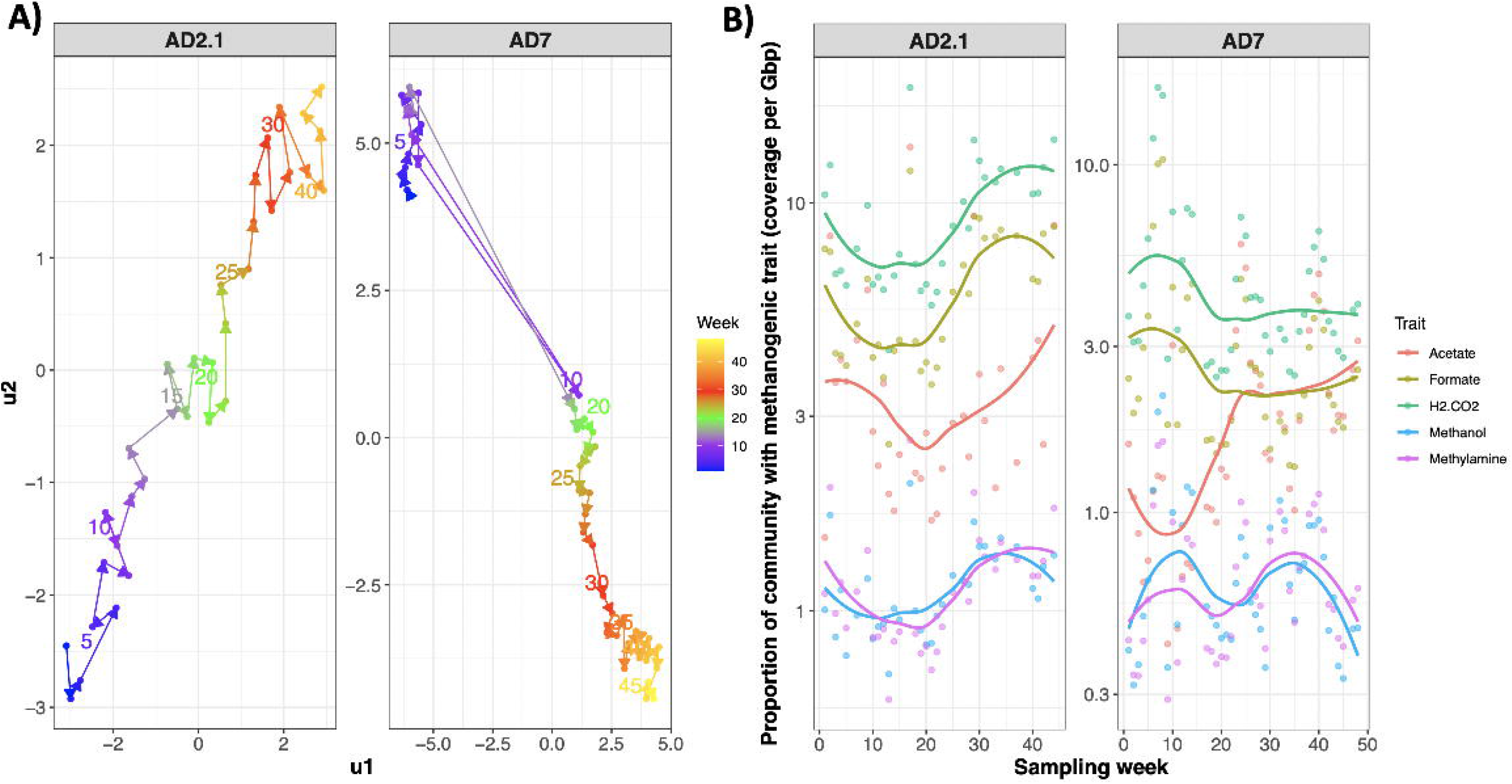
Archaeal dMAGs phylogeny and methanogens analysis. A phylogenetic tree created with all 45 Archaea dMAGs and representatives from GTDB, using a maximum likelihood approach (see *Methods*). Only the dMAGs are shown, along with their RED value encoded in the first coloured bar to the right of the tree (see legend). For methanogens, additional coloured bars show the total counts of the methanogen-specific *mcr* gene and the results of a machine-learning based prediction of their substrate specificity (see *Methods*). Also, for methanogens, the right most group of bars show their coverage across the 13 reactors. Branch colours indicate major classes, as shown in the legend.

Overall, there were 1910 dMAGs that were novel at the species level, expanding the currently available genomic information for major, as well as under-studied, phyla. We particularly note a 4.6-fold increase in the representation of DTU030 phylum in the GTDB, and almost a doubling of the OLB16 phylum. There were nine other bacterial phyla showing above 0.3-fold increase in representation (Fig. 1C). Additionally, there were 20 bacterial and 2 Archaeal dMAGs that represent novel, candidate classes (RED<0.48), and an additional 91 bacterial (with 6 CPR) ones that represent novel orders (RED<0.63). The two Archaeal dMAGs forming a novel class were within the Iainarchaeota phylum (Fig. 2), and a detailed analysis of their genomes are to be published in a separate study.

### The novel bacterial classes and orders comprise fermenters and syntrophs

On the *de novo* phylogenetic tree, 10 of the 20 bacterial dMAGs with a RED value below 0.48 formed three distinct sub-clades, consistent with them representing new classes (see Fig. S3-S5). One of these novel classes, comprising four dMAGs, is within the Firmicutes phylum and is most closely related to the Clostridia and Mahellia classes (Fig. S3A). In agreement with this phylogeny, KO-based clustering showed these four dMAGs to form a cluster, distinct from their closest phylogenetic neighbours (Fig. S3B). A machine learning algorithm, that we devised to infer functional traits from KO annotations (see *Methods*), indicated these dMAGs were capable of fermentation and arsenate reduction. Consistent with this, they contained KEGG modules associated with fermentation, including pyruvate to acetyl-CoA conversion and acetate production, as well histidine degradation. The second sub-clade, again comprising four dMAGs, is found within the Armatimonadota phylum, with half of its members classified as capable of fermentation and all of them carrying most of the genes for galactose degradation (Fig. S4). The third sub-clade comprises two dMAGs, most closely related to the classes Ignavibacteria and Bacteroidia within the Latescibacterota phylum. These dMAGs distinctively possess the metabolic modules for galactose, galacturonate and galactonate degradation and the associated Entner-Doudoroff pathway (Fig. S5). For each of these three novel classes, we picked a representative species, together with the AD reactor that it is found in, and have proposed a name, in accordance with the above findings and the latest naming guidance (*30*); *Candidatus Peptofermentans severntrenti* (AD20_Bin_2872), *Cand. Galactosivorans scrivelsbyensis* (AD18D_Bin_55) and *Cand. G. barfootsi* (AD2.2_Bin_614), and *Cand. Galactosutens copysgreenensis* (AD10_Bin_218).

Another seven dMAGs, with a mean RED value of 0.47, formed a distinct sub-clade, corresponding to a novel order within the Syntrophomonadia class (Fig. S6A). Members of this class are known syntrophs of methanogens, and accordingly, two of the dMAGs close to this sub-clade are assigned to *Syntrophomonas wolfei* (AD18H_Bin_253) and *Syntrophaceticus schinkiii* (AD2.1_Bin_491). Both of these syntrophic species have been co-cultured with hydrogen utilising methanogens (*31,32*) and are further identified in our interaction predictions (see below). We find that most of the dMAGs on this novel sub-clade harbour parts of the KEGG modules for glycolysis, pyruvate-to-acetylCoA oxidation, reductive acetylCoA (Wood-Ljungdahl) and acetate production pathways, lack the TCA module, and encode the leucine degradation pathways, and are classified as fermenters by our trait classifier (Fig. S6A). Additionally, six of the seven dMAGs form a distinct cluster based on their genome content, with their distinctive genes including a lyase (K01640) and a synthetase (K18661) involved in leucine degradation (Fig. S6B). Based on these findings, we hypothesize that these dMAGs specialise in peptide degradation and acetate production, possibly in association with acetate- or methylamine-utilising methanogens. In line with this hypothesis, we find additional evidence for interaction with methanogens (see below) for two of these novel dMAGs (MINI_Bin_251 and AD20_Bin_1939). Based on these findings, and the AD reactor from which they were found, we suggest the name *Cand. Polysyntrophus singletonbirchi* for one representative member (MINI_Bin_251) of this new order.

### Archaeal populations in individual AD communities are dominated by methanogens

AD communities are expected to have abundant methanogen populations, and we found representation of all known methanogen classes in our dataset (Fig. 2). In total, 38 different dMAGs were found (see Fig. 2), spanning five methanogen classes. For each methanogenic dMAG, we verified presence of the key methanogenesis marker gene *mcr* and used machine learning on overall KO content to infer their methanogenic substrate specificity (*see Methods* and Fig. 2). The most diverse class was the autotrophic Methanomicrobia (17 dMAGs), which oxidise hydrogen, followed by the Thermoplasmata (nine dMAGs), which had the highest association with methanol and methylamine as substrates (Fig. 2). These machine learning-based substrate assignments are broadly in agreement with culture-based studies of methanogenesis in representative species (*25, 33*). In addition to the next most diverse class, the Methanosarcinia (with seven dMAGs), we also found four Methanobacteria and just a single dMAG (AD2.2_Bin_1085) from the class Thermococci. This dMAG is assigned to the genus *Methanofastidiosum*, a newly described genus hypothesised to perform methylated thiol reduction (*34*). This dMAG is predicted by our KO-based approach to use hydrogen and methylamines as substrates for methanogenesis.

The overall abundance of the different methanogens formed a general trend that split reactors into two groups (see Fig. 2). The first group, comprised of 10 out of 13 reactors, was dominated by methanogens from the Methanosarcinia class, which harbours generalist methanogens (*25, 33*). Reactors AD2.1, AD2.2, AD7, AD10 and AD12 all shared the same most-abundant methanogen, a novel species (AD10_Bin_72) from the *Methanothrix* genus, members of which are known for their high affinity for acetate (*35*). AD20 was also dominated by a *Methanothrix* species. Reactors AD16.1, AD16.2, AD18D and MINI were dominated by a single methanogen, a *Methanosarcina flavescens* (AD2.1_Bin_512), that is shown to be a generalist methanogen able to grow on hydrogen, acetate, or methylamines (*36*), as also observed in our trait assignment (Fig. 2). The second group consists of three reactors, AD3, AD11 and AD18H, which are dominated by methanogens from the Methanomicrobia and Methanobacteria classes. Notwithstanding these dominant dMAGs in specific reactors, we find significant shifts in the abundance of different methanogen types over time in most reactors, as discussed below.

### Longitudinal dMAG coverage analysis identifies interaction partners of methanogens

As discussed above, AD communities are expected to be strongly structured by pairwise mutualistic syntrophic interactions in which syntrophic partners produce acetate, hydrogen, formate, methanol, or methylamines that are utilised by the different groups of methanogens (*25*). To identify possible syntrophic associations of these diverse methanogens, we searched for non-methanogenic dMAGs that showed highly correlated coverage profiles with specific methanogens across individual reactor time series (see *Methods*). We focused on those pairs where one species was a methanogen (38 methanogen dMAGs) and the other a possible syntroph, either bacterial or archaeal (2202 ‘partner’ dMAGs). We allowed each potential syntroph to be associated with only one methanogen, but a methanogen could have more than one syntrophic partner.

This analysis identified 518 possible interactions (with a correlation coefficient above 0.9) among 450 possible syntrophs and 33 methanogens, since some interactions involved the same partner in multiple reactors (see Fig. 3 for example associations). Many of the identified pairs matched well-established and unambiguous syntrophic interactions from the literature, for instance the species *S. wolfei* and *S. schinkii* mentioned above, *Syntrophus aciditrophicus* (AD7_Bin_431), *Flexilinea floccule* (AD20_Bin_801), and *Coprothermobacter proteolyticus* (AD20_Bin_2104). The first three species are model organisms for a syntrophic lifestyle, with two of them discussed above (see Fig. 3A/B for *S. wolfei* case). The *S. aciditrophicus* is a fatty-acid-degrading specialist that grows in syntrophy with hydrogen-utilising methanogens (*37,38*). In our analysis, it is predicted to interact with a methanogen (AD20_Bin_1426) from the class Thermoplasmata (Fig. 3C/D), members of which are shown to require hydrogen for growth (*39*, see also below). The *C. proteolyticus* is shown to specialise in degradation of proteinaceous substrates in syntrophy with *Methanothermobacter thermautotrophicus* (*40*). *F. floccule* is a fermentative anaerobe isolated from anaerobic sludge granules and grows best in co-culture with a hydrogenotrophic methanogen (*41*). In our analysis, both of these species are predicted to interact with dMAGs (AD18H_Bin_493 and AD2.1_Bin_909, respectively) assigned to the Methanobacteria class and that are predicted to use hydrogen (Fig. 2).

**Figure 3.**
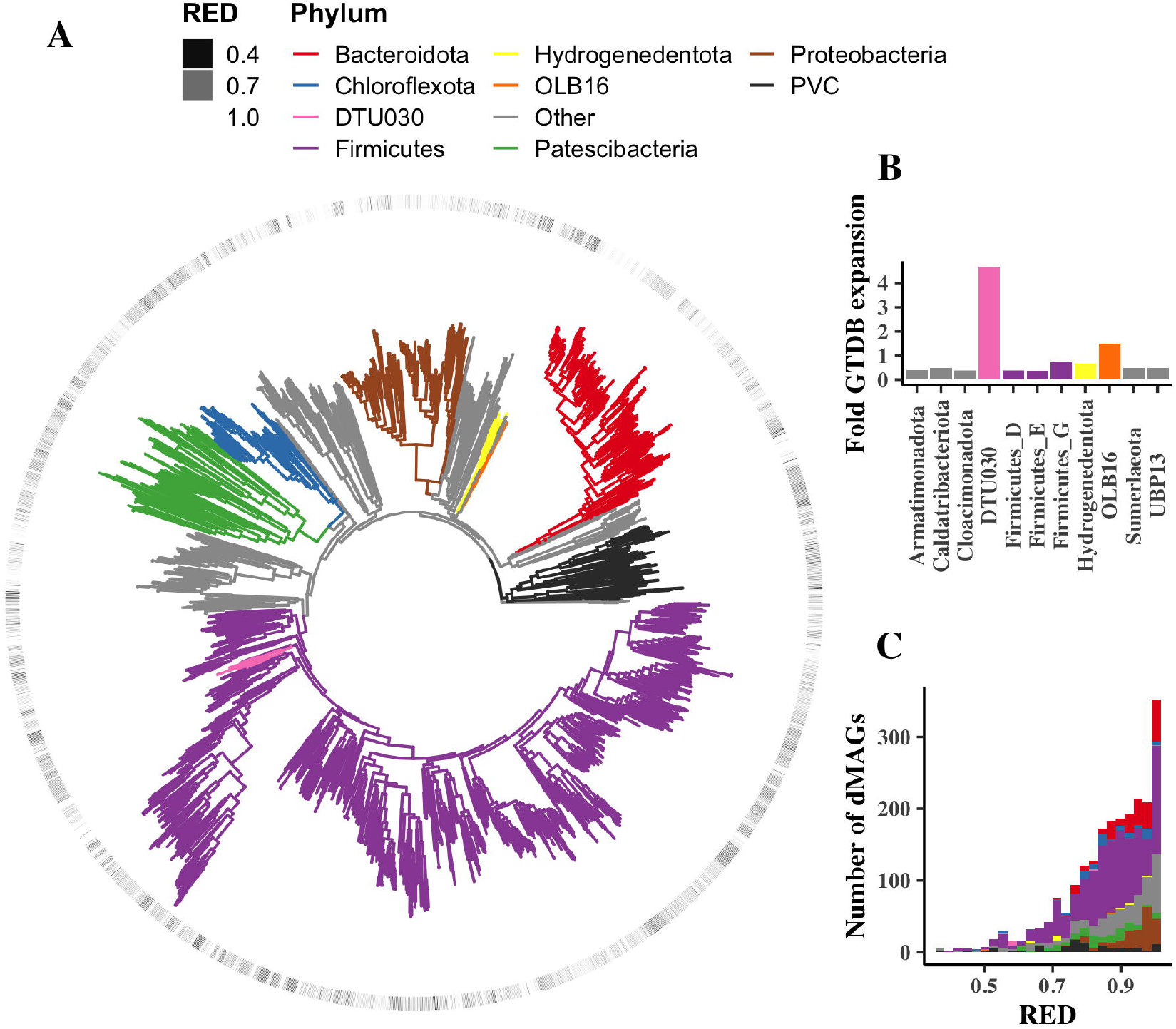
Examples of strong positive correlations found in AD reactor communities. **(A)** Time series of dMAG normalised coverage depths for MINI_Bin_333 (*Methanosarcina* sp002509325 - blue) and AD18H_Bin_253 (*Syntrophomonas wolfei -* red) in reactor AD18H. **(B)** Pairwise correlation of log-transformed normalised coverages for MINI_Bin_333 and AD18H_Bin_253 (r = 0.98, p = 8.50e-13). **(C)** Time series in reactor AD20 of AD20_Bin_1426 (*Methanomethylophilaceae sp*. – blue) and AD7_Bin_431 (a known syntroph, *Syntrophus aciditrophicus* - red). **(D)** Pairwise correlations of log-transformed normalised coverages for AD20_Bin_1426 and AD7_Bin_431 (r = 0.95, p = 3.69e-16).

Multiple further lines of evidence support the idea that predicted associations reflect real metabolic interactions. Firstly, there is a strong taxonomic signal to the potential interactions. The different methanogen classes differ greatly in the frequency with which they form associations. The highest number of associations is found in the Methanobacteria, with 36.75 associations, on average, per dMAG, and they are also very frequent in the Thermococci (27) and Thermoplasmata (16.34), but less so for the Methanomicrobia (6.82) and Methanosarcinia (11.57) (see also Fig. 2). Considering the taxa of both partners, we obtain a contingency table that strongly indicates a non-random association of the partner’s taxonomic phylum with the particular methanogen classes (p-value = 6.829e-10, Chi-squared test) (Fig. S7). Interestingly, we found that methanogens from Thermococci, Thermoplasmata and Methanomicrobia all had a high probability of correlations with Bathyarchaeia from the phylum Crenarchaeota (Fig. S7). Specific examples of these interactions are discussed further below from a metabolic perspective. In addition, a large number of interactions were found between methanogens from the Methanosarcina class and the phylum Elusimicrobia, members of which are found in termite and human guts, as well as a diversity of other environments, and are associated with acetate and H_2_ production (*42*).

Secondly, the KEGG ortholog gene profiles of the non-methanogenic partner differed depending on the methanogen with which it correlated (permutation ANOVA on KEGG assignments - R2 = 0.10122 p < 0.001 for methanogens with at least five partners). This implies a degree of metabolic selection in the non-methanogenic partner. Thirdly, we compared the genomic content, in terms of KEGG orthologs, between the 450 partner dMAGs (Pearson’s r>0.9 with a methanogen) and 457 non-partner dMAGs (no Pearson’s r > 0.5 with any methanogen in any reactor – see *Methods*, Fig. S8 and Table S2). Overall, we found significant differences in functional gene composition between these two groups (2764 genes with false discovery rate < 0.01). The most significant gene with a large frequency difference between the two groups was the gene K04069, which encodes for ‘pyruvate formate lyase activating enzyme’ – a protein which activates the enzyme pyruvate formate lyase, which catalyses the conversion of pyruvate and coenzyme-A (CoA) to formate and acetyl-CoA. This is consistent with the important role of formate in methanogenic syntrophies (*43,44*). Furthermore, several of the genes enriched in the partner group are, or are associated with, oxidoreductases, which have previously been found to be important in syntrophic interactions (*24, 45, 46*), as well as genes involved in sugar transport and metabolism, indicating fermentative metabolism in the partners (see Fig. S8 and Table S2). To further see if these differences in gene content are predictive, we used a random forest algorithm (see *Methods* – Table S3) to attempt to distinguish, from their KEGG ortholog frequencies alone, whether a dMAG was a methanogen partner. This approach achieved an overall accuracy of 83.46%, indicating there is a strong genomic signal associated with being a methanogenic partner or not.

The majority of correlating dMAG pairs, 309 out of 510, occur together in more than one reactor. However, in only 7 of these 309 cases, do we find that the dMAG pairs strongly correlate in more than one reactor. Two of these pairs involved the same generalist methanogen from the Methanosarcinia class (AD2.1_Bin_512), while the bacteria involved (AD2.2_Bin_49) were from the Endomicrobia class (Elusimicrobia phylum), and from the Clostridia class within the Firmicutes phylum (AD2.1_Bin_784). Based on the observation that not all highly-correlating pairs are found to correlate in all reactors, we hypothesise that at the dMAG – i.e. approximate sub-species – level, syntrophies are not obligate, but that the interaction develops only between some strains in some reactors, possibly as the result of coevolution. In support of this hypothesis, we find that in general there is a high level of genome variability between MAGs comprising a given dMAG (mean variance in KOs across MAGs within a dMAG is 6.28%). In Fig. S9 we give as an example, the methanogenic Thermococci (AD2.2_Bin_1085, *Methanofastidiosum*), and the Bathyarchaeia (MINI_Bin_276). This pair cooccurs in five reactors, and in one (AD7) we flag them as potential syntrophs (r = 0.95, p-value = 4.59e-22) and in another (AD20) they may be associated (r = 0.72, p-value = 3.2e-6) although below our strict threshold, but in the three others (AD2.1, AD2.2 and AD12) they have no significant association (p > 0.1). However, the KEGG ortholog profiles for the seven MAGs assigned to the *Methanofastidiosum* dMAG AD2.2_Bin_1085 do vary, differing at a median of 1.9% of KOs between strains (i.e MAGs). Intriguingly, the MAG with the most similar KO profile to the strain in AD7, where the correlation is observed, is in fact the AD20 dMAG (which is only 1.3% divergent). The Bathyarchaeia had a somewhat higher level of genomic variability amongst its five MAGs differing at a median of 6.4% of KOs. In this case we cannot test if the strain in AD20 is most similar to the AD7 strain as this dMAG was not binned in AD20 but its coverage determined through mapping (see *Methods*).

### Predicted interactions highlight an association of Bathyarchaeia and Patescibacteria with methanogenesis from methylated compounds

The above analyses suggest that predicted interactions represent testable hypotheses for understanding methanogen interactions in AD communities, either syntrophic or otherwise. Among specific predicted interactions, we highlight the strong correlation of the single Thermoccoci methanogen and several Thermoplasmata methanogens with Bathyarchaeia in our dataset (Fig. S7). In the case of Thermococci (AD2.2_Bin_1085, *Methanofastidiosum*), the interaction is with the Bathyarchaeia (MINI_Bin_276), as mentioned above. Analysis of KO content of these dMAGs shows that the *Methanofastidiosum* encodes for a single methyltransferase and otherwise lacks the genes for the upper part of the methanogenesis pathway, as seen in all of the Thermoplasmata methanogens (Fig. S10). This is indicative of hydrogen or acetate dependent methanogenesis from methylated substrates, as predicted before from other metagenomics studies for *Methanofastidiosum* and experimentally shown for several Thermoplasmata species (*39, 47*). The *Methanofastidiosum* encodes for the *frh* gene encoding a F420-linked hydrogenase, which is not found in any of the Thermoplasmata dMAGs, and the acetyl-CoA synthetase gene that converts acetate to acetyl-coA and that is found only in few of the Thermoplasmata dMAGs. These findings suggest that the *Methanofastidiosum* in our system is capable of both H_2_- and acetate-based syntrophies. All of the Bathyarchaeia dMAGs encode the acetyl-CoA synthetase gene and the acetyl-CoA decarboxylase/synthase complex (ACS), as well as the associated carbon monoxide dehydrogenase (Fig. S10), indicative of capacity for acetogenesis. Unlike a previous analysis (*48*), but similar to a more recent one (*49*), we find that none of the Bathyarchaeia dMAGs encode the genes of the *mcr* complex. This gene-based analysis is thus supportive of an acetate- or hydrogen-dependent syntrophy between the *Methanofastidiosum* and Bathyarchaeia, that could explain the predicted partnership. The Bathyarchaeia have been predicted to undertake either reductive acetogenesis or syntrophic acetate oxidation with H_2_ production in recent metagenomic analyses (*48, 49*), but a possible interaction with methanogens, as predicted here, is to be tested.

As mentioned above, we find that six out of nine Thermoplasmata methanogens in our dataset have strong partnerships also with Bathyarchaeia, and with Patesciebacteria (see Fig. S7). Considering that all analysed Thermoplasmata species to date have shown a hydrogen- or acetate-dependent methanogenesis from methylated substances (*39, 47*) and are implied to have a competitive advantage over hydrogenotrophic methanogens in hydrogen utilisation (*50*), these partnerships could again involve a hydrogen-based syntrophy between these methanogens and acetogenic Bathyarchaeia. While members of the Patescibacteria have frequently been suggested to be involved in symbiotic associations due to their reduced genomes (*4, 12*), an association with methanogens, as implied by our analysis, has not been suggested so far and needs to be tested. To this end, we find a significant enrichment of 10 specific KEGG orthologs in Patescibacteria (17 dMAGs) interacting with Thermoplasmata, compared to other Patesciebacteria in our datasets (110 dMAGs) (see Table S4). These genes do not include the oxidoreductases characteristic of the general dMAG syntrophs, mentioned above, suggesting other mechanisms for these putative syntrophies.

### Reactors exhibit dramatic shifts in community structure and methanogenesis pathways

The comprehensive nature of our dMAG collection, comprising 41 million reads per sample, enabled us to use the dMAG normalised abundances to profile both community diversity and structure over time. The total number of dMAGs observed in each reactor remained mostly constant over time but did differ significantly between reactors (Fig. S11); however, detecting the definitive absence of a dMAG is difficult giving sampling issues. We therefore also examined the dMAG diversity as measured by the Shannon diversity, which is more robust to low abundance dMAGs. This did show some trends, decreasing significantly over time in AD11 (p = 2.1e-08) and AD20 (p = 1.0e-3) but increasing in AD7 (p = 4.8e-05) (Fig. S12). In contrast, the community structure as measured by the full dMAG abundance profile exhibited clear shifts with sample time for all reactors. In Figure 4A we give examples of such community structure shifts for two reactors, AD2.1 and AD7, using the umap dimension reduction algorithm to plot the 2240-dimensional vectors of weekly dMAG coverages in two dimensions (see *Methods*). Similar changes were observed across all the reactors (see Fig. S13). We used multivariate permutation ANOVA of community structure against sampling week to verify that these shifts were statistically significant for all reactors (see Table S5) - for AD20, changes were small but still of marginal significance. For a number of reactors these shifts appear cyclic (notably AD12 and AD20, Fig. S13). These changes in community structure could not be entirely explained in terms of changes in feed or operating conditions. Operating conditions did impact community structure but when we included both operating conditions (e.g. pH, temperature and oxygen) and the proportions of the different feed types, along with sampling week, in the ANOVAs, the sampling week still explained more variation (between 20.2% and 49.3%) than any other variable for every reactor for which we had sufficient metadata to test (see Table S6 and Fig. S14).

**Figure 4.**
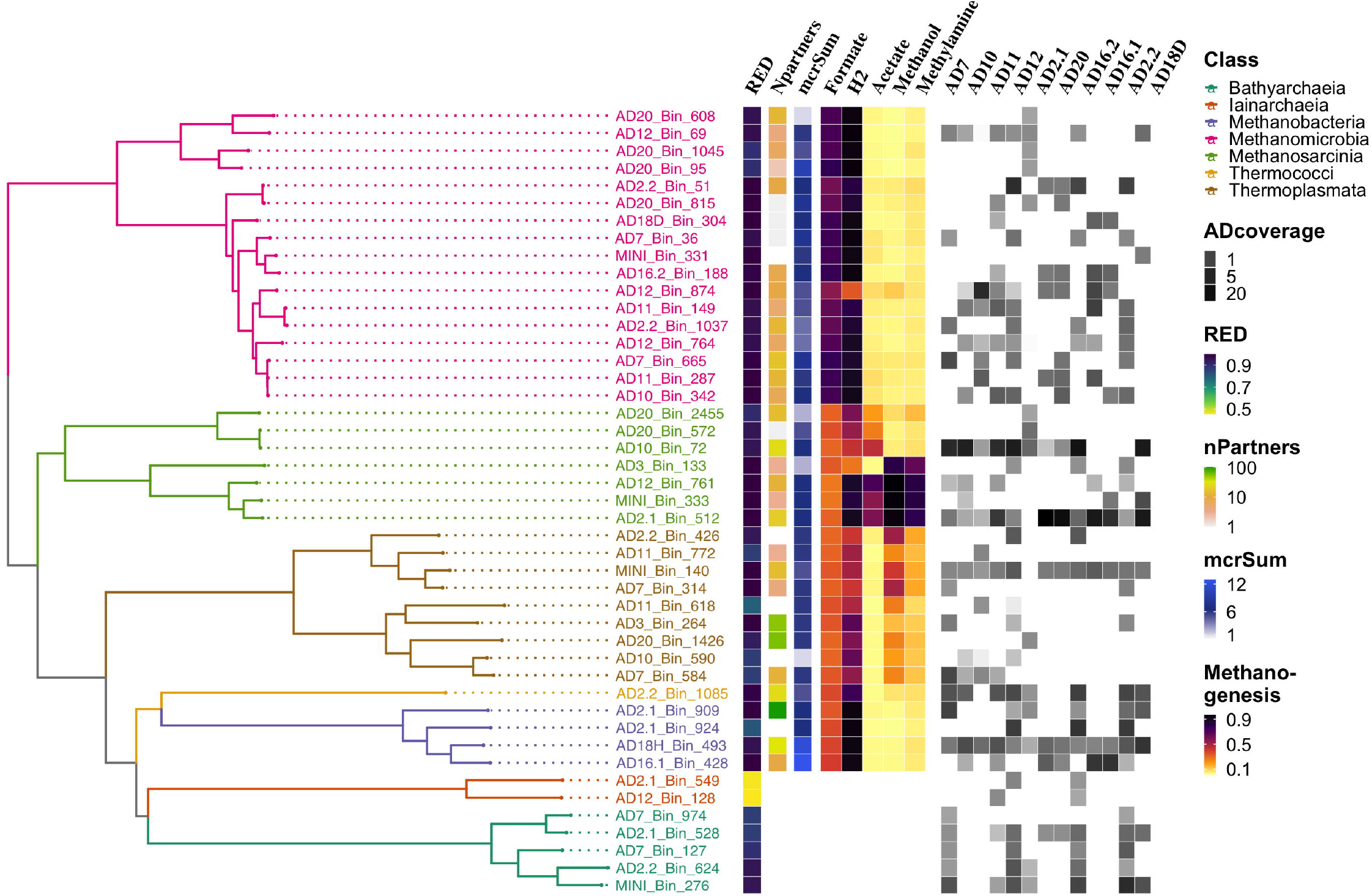
Changes in community structure and methanogenesis pathways over time for two reactors AD2.1 and AD7. **(A)** Time series of dMAG normalised abundances mapped to two dimensions using umap (multivariate ANOVA R2 = 0.355 p < 0.01 and R2 = 0.418 p < 0.001 for AD2.1 and AD7 respectively). Individual data points are colored according to sampling time and as indicated on the legend. **B)** Time series of total normalised coverage of dMAGs capable of utilising different substrates for methanogenesis (multivariate ANOVA R2 = 0.1730 p = 0.012 and R2 = 0.1030 p = 0.014 for AD2.1 and AD7 respectively). Each group of dMAGs with a specific methanogenic trait (i.e. pathway) is shown with a colored line as indicated on the legend.

For most reactors, these changes in overall community structure were associated with a change in the overall functional capacity to utilise different methanogenic substrates (Fig. 4B). We used the methanogenic substrate assignments for the dMAGs discussed above to calculate the proportion of the community (in terms of normalised coverage) capable of performing methanogenesis with a particular substrate (see *Methods*). Multivariate ANOVA revealed that for eight out of 12 reactors the methanogenic pathways shifted significantly over time (see Table S7). In Fig. S15 we plot the methanogenesis trait abundances over time for all eight of these reactors – except for AD18H and AD18D, which had incomplete time series. This reveals that in some reactors there is a change in relative importance of different types of methanogenesis, e.g. around week 20 in AD7 acetoclastic methanogenesis becomes more important (see Fig. 4B), whilst other transitions involve an increase in the abundance of all methanogen types, e.g. in AD16.1 (see Fig. S15).

### Particular dMAGs correlate with community productivity in terms of methane yield accounting for operating temperature and reactor identity

A key measure of community productivity for AD reactor communities is the methane yield, which we define here as the total volume of methane gas produced per unit mass of feedstock. This is shown as a function of time for all reactors where we were able to gather these data in Fig. S16. To provide more robust methane yield statistics, we filtered outlying values and removed reactors that were paired or multi-stage (see *Methods*). Since different reactors have different feedstocks this means that yield cannot be compared across reactors although changes within a reactor over time are valid if we assume that substrate COD (chemical oxygen demand) for a particular reactor is not changing substantially. We do see substantial fluctuations in yield for most reactors over time in Fig. S16. To determine what drives these fluctuations we performed multivariate regressions of yield against operating conditions but with reactor as an additional quantitative variable to account for differences in potential feedstock COD and reactor design. The strongest relationship was found with operating temperature which had a significant positive impact on yield (regression coefficient 15.297, p = 0.003; see Table S8).

To determine any additional impact of community structure on performance, we included both reactor identity and temperature in multivariate regressions of yield against individual normalised log transformed dMAG abundances. This strategy should find dMAGs that are important to yield across multiple reactors, while accounting for mean performance differences between reactors and changes in operating temperature. We only tested dMAGs that were present in at least half of all samples to focus on trends that can be consistently identified across reactors. This revealed multiple dMAGs with correlations to performance (300 out of 588 tested with adjusted p-value < 0.05 see Table S9 and Fig. S17). These dMAGs included the novel methanogen from the class Thermococci (AD2.2_Bin_1085) discussed above. The dMAGs highly significantly positively associated with yield (adjusted p-value < 0.001) were also more likely to be potential syntrophic partners to methanogens as defined above on the basis of coverage profile correlations. The proportion of syntrophs amongst these dMAGs was 24.4% vs. 11.9% for the rest of the dMAGs tested (see Table S10 Fisher’s exact test p-value = 0.003).

Notwithstanding these findings, however, we note that the community structure had a relatively small impact on methane yield beyond that explained by reactor identity and temperature alone. The total variance explained in a random forest model including the significant dMAGs, reactor identity, and temperature was 90.48% versus 88% for a multivariate regression of yield against just the reactor identity and operating temperature.

## CONCLUSION

We presented here analysis of a comprehensive longitudinal sampling of microbial communities coupled to genome-resolved metagenomics. By focusing this analysis on industrial AD reactors we were able to combine metagenomics with metadata from the same reactors, and use these integrated data to identify novel species, alonng with their interactions and their contribution to community function. The resulting information proves that longitudinal metagenome analyses, such as this one, can become an essential tool for understanding microbial communities and predicting their dynamics. In the former direction, we believe that the high number of strongly positive correlations identified from the coverage profiles (i.e. predicted interaction partners) provides evidence of the highly structured nature of AD communities and constitutes a useful set of experimentally testable hypotheses for putative ecological associations. The fact that we could reliably (83.46% accuracy) predict which microbes correlated with methanogens from their genomes alone gives us confidence that these reflect real interactions. In particular, future experimental studies using culture enrichment techniques could be used to verify the predicted associations of Bathyarchaeia and Patescibacteria with Thermocci and Thermoplasmata from our data.

We note, however, that our results provide an alternative model for syntrophic associations in anaerobic communities. We propose that these are more facultative in nature than hitherto appreciated, with the exact strain of microbes being important as to whether a potential syntrophy is realised. It is possible therefore that isolates that are syntrophic when co-cultured may not actually be interacting in the community from which they are taken, and a degree of co-evolution might be necessary for these associations to be realised, as observed in some syntrophic co-culture studies (*46, 51*). This emphasizes the importance of using strain-resolved metagenomics approaches, as demonstrated here.

Our analysis reveals significant structural changes in AD communities over time associated with shifts in the importance of different types of methanogenesis. Shifts in community structure have been observed in other studies, that used temporal data, despite having much lower resolution and focusing on only a few, or two (start – end), time points (*17, 20, 26*). The higher resolution of the data presented here indicates possible cyclic dynamics in community structure, which needs to be further examined. We presented here, for the first time, a trait-based approach to associate these shifts to potential major changes in community metabolic function. This could be confirmed in future, with methods such as transcriptomics or proteomics to verify that this change in potential is related to realised function. Notwithstanding such shifts in community structure and methanogenesis pathways, we find that a main determinant of methanogenic community performance is temperature. In this context, we note community structure was shown to change significantly when cold-adapted methanogenic communities were subjected to temperature increase (*52*), or when AD reactor communities were adapted to a new feed (*53*).

Further understanding of intrinsic dynamics of community structure, as well as its responses to external factors can benefit from longitudinal studies of integrated metagenomics and metadata, as presented here. It is only through such studies that we can achieve development of mechanistic or statistical models of community function, stability, and performance that have true predictive capability.

## METHODS

### Sampling information

We sampled DNA from 13 industrial-scale AD reactors for approximately a year on a weekly basis and recorded their operational parameters and methane productivity, but not every week was available for every reactor. Table S1 lists key parameters of these reactors along with number of sampling weeks for each. For all reactors, we had over 30 sampling times, with a maximum of 39 weeks. All reactors except one were fed with vegetable waste supplemented with animal manure and were operated around 40°C (see Table S1). One reactor, labelled AD20, was part of a sewage treatment plant, while another two reactors, labelled AD18.D and AD18.H, were connected in series as hydrolysing and digesting systems kept at 48°C and 43°C respectively. While different metadata were made available for each reactor, it was possible for all reactors to tabulate temperature, pH, methane concentration, hydrogen sulphide concentration, gas production rate, feed rate and feed type. Biological samples were taken directly from AD reactors by facility operators on a weekly basis and shipped in ice-cooled containers. Upon receipt, they were first kept in a 4°C fridge and then sampled into several 1-5ml aliquots within days. Most aliquots were processed within days for DNA extraction, while some were first stored in a -80°C freezer until subsequent thawing and extraction. DNA extraction was done using Qiagen DNeasy PowerSoil kit (Cat No./ID: 12888-100) following the manufacturer’s manual.

### Data and code availability

All of the 2240 dMAGs built from this work are deposited on the ENA as part of project PRJEB39861. The pipeline for binning is freely available from https://github.com/Sebastien-Raguideau/Metahood and that for the trait inference described below at https://github.com/chrisquince/genephene.

### Sequencing

Sequencing was carried out in two-step process starting first by set of 15 pilots samples sequenced at 2×250. This was followed by a total of 375 DNA libraries pooled together and sequenced over six NovaSeq S2 lanes and one S4 lane at 2×150. This yielded an average of 41M reads per sample.

### Sequence analysis and binning

Reads were quality filtered and adaptors removed using Trimgalore version 0.6.4_dev (*54*), a wrapper around cutadapt (*55*) and fastqc (*56*). A total of 13 reactor-wise co-assemblies were carried out using megahit (*57*) version v1.2.9. Samples were mapped to assemblies using bwa_mem version 0.7.17-r1188 (*58*). The resulting bam files were handled using samtools v1.5 (*59*) and bedtools v2.28.0 (*60*) to produce mean contig coverage. Each reactor coassembly was binned using Concoct version 1.1.0 (*61*).

### Bin assessment

For all bins from the 13 assemblies, gene prediction was carried out using prodigal V2.6.3 (*62*), with option “-p meta”. A selection of 36 single-copy core genes (SCGs) were annotated through rpsblast v2.9.0 (*63*) using the pssm formatted COG database (*64*), which is made available by the CDD (*65*). Only bins with at least 50% of the SCGs in a single copy where kept. The resulting 7240 bins were dereplicated using drep v2.6.2 (*68*) with default settings: alignment of at least 99% similarity for any section of the genome spanning a minimum of 10% of bin length. This resulted in 3375 bins. These bins were taxonomically annotated using GDTB-Tk v2.6.2 (*27*) with GTDB v95 (*28*). Bin quality was assessed with CheckM v1.1.3 (*66*), using custom marker sets as follows: bins classified as bacteria, other than Patescibacteria, and archaea were assessed using respective marker sets from GToTree (*67*), while bins classified as Patescibacteria were assessed using specific marker set made available for CheckM (*3*). For all 3375 bins from drep, the represenetative bins is chosen as the ones that have the maximum value for “completion-contamination”. Any bins with a “completion-contamination” score over 75% are considered a MAG. Those MAGs that are a cluster representative are called a dMAG (i.e dereplicated MAG). In total, 5303 MAGs and a corresponding 2240 dMAGs were obtained.

### Coverage analysis

Each of the 5303 MAGs are uniquely related to one of the 13 co-assemblies and their corresponding samples. We could therefore assign an unambiguous coverage to a dMAG when a MAG from that dMAG was present in a reactor (Fig. S1). Mean coverage of a MAG was obtained by taking the sum of nucleotides mapping to all its contigs and dividing by total length of the MAG. The strict parameters used for MAG dereplication ensured that each of the 2240 dMAG clusters possess only one MAG in each reactor co-assembly. This will miss dMAGs that were present in a reactor but failed to be binned. We therefore mapped contigs from the co-assemblies to the dMAG collection. A dMAG is considered present in a reactor, if more than 10% of the dMAG could be found with identity > 99%, and its coverage profile over corresponding samples is computed as an average over the mapped contigs(Fig. S1). If a dMAG had no binned representative in a reactor and its contigs failed to recruit from the co-assembly then its coverage is set to zero for all samples from that reactor. Coverage values were normalised using the total read length from a sample and divided by 1.0e9 to give coverage depths per Gbp of sequence.

### Phylogenetic analyses of dMAGs

*De novo* phylogenetic trees were built using FastTree v2.1.10 (*69*) using previously generated GDTB-tk marker set. Tree visualisations were created with custom R scripts using the ggtree library (*70*).

### Functional analyses of dMAGs

Annotation of KEGG orthologs (KOs) (*71*) for all dMAGs was obtained using kofamscan v1.2.0 (*72*). The completeness of each KEGG module in the dMAGs was tabulated using the R library METQy (*73*).

### Cluster analysis of dMAGs

Where utilised, clustering of dMAGs was performed using their KO annotations. Clustering was done using the kmeans method, as implemented in the “fviz_nbclust” function of the R package factoextra.

### Functional trait inference for dMAGs

We used a machine learning approach to assign functional traits to dMAGs (*74*). Two trait databases were used, FaproTax which contains a set of generic functional and metabolic traits relevant to environmental microbes mapped to microbial species, and a dedicated methanogen database that included growth information on the methanogenesis substrates: acetate, H2 and C02, formate, methanol, and methylamine (*75*). In both cases models were constructed to predict traits using as an input KEGG ortholog frequencies obtained by mapping the database taxa to RefSeq genomes and annotating against the KEGG database. Models were validated using ten-fold cross-validation. For the FaproTax traits a logistic regression with a Lasso penalisation was found to perform best, but for the methanogensis substrate prediction, a support vector machine was used. The FaproTax models were applied to all dMAGs to provide a summary of functional information, while the methanogenesis substrate prediction was applied only to those archaeal dMAGs that had at least one copy of the *mcr* gene. Details of the modelling methodology are available in (*74*). In brief, to calculate trait abundances at the sample level, a matrix multiplication of the dMAG trait assignments with the dMAG normalised coverage matrix was performed to sum the coverage of all dMAGs with a particular trait in a sample.

### Time series analyses

To identify possible syntrophic associations, we searched for pairs of dMAGs that were highly correlated across individual reactor time series (Pearson’s correlation r > 0.90 and Benjamini-Hochberg adjusted p-value < 1.0e-5) based on log-transformed normalised coverage. We focused this analysis on those pairs, where one of the pair was a methanogen and the other a possibly syntrophic partner, either bacterial or archaeal. We allowed each non-methanogen to be associated with only one methanogen, but a methanogen could have more than one partner. For comparison to these dMAGs we also defined non-syntrophic partners as those dMAGs that do not correlate with any methanogens with r > 0.5 in all reactors.

## Acknowledgements

We would like to thank Dr. Mary Coates for project coordination and Dr. Mark Pallen for help with species naming. We acknowledge the following individuals and companies for providing samples; Alex Semenyuk (Barfoot Energy Ltd.), Dr. Stephen Temple (J.F. Temple & Son Ltd.), Chris Meyer (Singleton Birch Ltd.), Dr. Ed Moorhouse (Shropshire Energy UK Ltd.), Henry Dymoke (Scrivelsby Biomass Ltd.), and Peter Vale (Seven Trent Water Ltd.).

## Notes

**Funding key:** This project is funded by the Biotechnology and Biological Sciences Research Council (BBSRC) grants (BB/N023285/1 and BB/K003240/1). CQ is funded through MRC MR/S037195/1 and the CLIMB-BIG-DATA consortium MR/T030062/1. GC is funded by Science Foundation Ireland through a Career Development Award (17/CDA/4658). OSS acknowledges additional funding from Gordon and Betty Moore Foundation (Grant GBMF9200, https://doi.org/10.37807/GBMF9200).

### Competing Interest Statement

The authors have declared no competing interest.

