## Supplementary Figures and Tables for "Novel microbial syntrophies identified by longitudinal metagenomics"

This supplementary information comprises 10 supplementary tables and 16 supplementary figures, as included below.

### SUPPLEMENTARY TABLES

| Reactor ID | Company ID | Temporal coverage (weeks) | Feedstock type and mean daily quantity (approx.) | Operating Temperature (˚C) |
|---|---|---|---|---|
| AD7 | C | 44 | Cattle slurry, maize silage, beet, whey. 12T/day | 42 |
| AD10 | D | 36 | Pig manure, beet, maize, grass. 100T/day | 40 |
| AD11 | D | 35 | Maize, chicken litter. 55T/day | 40 |
| AD12 | D | 34 | Maize, beet, pig manure. 28T/day | 40 |
| AD2.1 | A | 33 | Maize/corn silage, rye grain, vegetable liquid food waste. 60 T/day | 39.5 |
| AD20 | G | 32 | Sewage waste | 38 |
| AD16.2 | E | 30 | Maize, rye, onions, chicken litter. 131T/day | 35-40 |
| AD16.1 | E | 26 | Maize , rye , onions , chicken litter. 131T/day | 35-40 |
| AD2.2 | A | 26 | Maize/corn silage, rye grain, veg., food waste. 60 T/day | 39.5 |
| AD18.D | F | 21 | Maize , grass , straw , chicken litter , cattle manure. 27T/day | 43 |
| AD18.H | F | 19 | Maize , grass , straw , chicken litter , cattle manure. 27T/day | 48 |
| AD3 | B | 14 | Maize/rye silage ~ | 41.7 |
| MINI | D | 8 | Maize, Straw. 0.002T/day. | 40 |

**Supplementary Table 1. List of AD reactors sampled for this study.** Summary of industrial reactors sampled for this study and their feedstock type. Reactors were sampled for a 50-week period, however, coverage was not perfect for any reactor with each one missing some weeks of data as indicated. All reactors were operated at 35-42˚C and with gas-based or mechanical stirring.

| KO id | KO name | p | q | tau | Mean non-partners | Mean partners | Diff. |
|---|---|---|---|---|---|---|---|
| K04069 | pyruvate formate lyase activating enzyme | 5.68E-36 | 2.12E-32 | 157 | 1.12 | 2.33 | 1.2 |
| K02355 | elongation factor G | 1.83E-35 | 3.42E-32 | 154 | 1.32 | 1.99 | 0.674 |
| K07037 | cyclic-di-AMP phosphodiesterase PgpH | 1.55E-34 | 2.32E-31 | 150 | 0.319 | 0.753 | 0.434 |
| K18682 | ribonucrease Y | 8.96E-34 | 1.11E-30 | 147 | 0.584 | 0.964 | 0.38 |
| K06881 | bifunctional oligoribonuclease and PAP phosphatase NrnA | 5.14E-33 | 5.48E-30 | 143 | 0.499 | 0.913 | 0.414 |
| K01493 | dCMP deaminase | 2.63E-31 | 1.63E-28 | 135 | 0.499 | 0.896 | 0.397 |
| K07238 | zinc transporter, ZIP family | 1.38E-29 | 6.07E-27 | 128 | 0.54 | 1.09 | 0.553 |
| K06960 | uncharacterized protein | 4.51E-27 | 9.9E-25 | 116 | 0.44 | 0.816 | 0.376 |
| K00282 | glycine dehydrogenase subunit 1 | 5.32E-27 | 1.13E-24 | 116 | 0.206 | 0.562 | 0.357 |
| K13993 | HSP20 family protein | 2.71E-26 | 5.33E-24 | 113 | 0.792 | 1.52 | 0.723 |
| K00335 | NADH-quinone oxidoreductase subunit F | 6.67E-25 | 9.96E-23 | 106 | 0.681 | 1.52 | 0.837 |
| K18672 | diadenylate cyclase | 1.3E-24 | 1.84E-22 | 105 | 0.532 | 0.902 | 0.37 |
| K01843 | lysine 2,3-aminomutase | 4.09E-24 | 5.09E-22 | 103 | 0.188 | 0.542 | 0.354 |
| K01006 | pyruvate, orthophosphate dikinase | 5.73E-24 | 6.69E-22 | 102 | 0.663 | 1.08 | 0.419 |
| K07456 | DNA mismatch repair protein MutS2 | 4.57E-23 | 4.27E-21 | 97.8 | 0.479 | 0.853 | 0.374 |
| K07079 | uncharacterized protein | 7E-23 | 6.37E-21 | 97 | 0.427 | 0.967 | 0.54 |
| K00176 | 2-oxoglutarate ferredoxin oxidoreductase subunit delta | 3.34E-21 | 2.24E-19 | 89.3 | 0.352 | 0.811 | 0.459 |
| K01619 | deoxyribose-phosphate aldolase | 1.26E-19 | 6.26E-18 | 82.1 | 0.606 | 0.967 | 0.361 |
| K18332 | NADP-reducing hydrogenase subunit HndD | 1.22E-18 | 5.13E-17 | 77.7 | 0.206 | 0.564 | 0.359 |
| K00334 | NADH-quinone oxidoreductase subunit E | 3.55E-18 | 1.36E-16 | 75.6 | 0.716 | 1.41 | 0.693 |
| K01926 | redox-sensing transcriptional repressor | 5.13E-18 | 1.92E-16 | 74.8 | 0.42 | 0.827 | 0.407 |
| K00177 | 2-oxoglutarate ferredoxin oxidoreductase subunit gamma | 9.72E-18 | 3.54E-16 | 73.6 | 0.409 | 0.844 | 0.435 |
| K04486 | histidinol-phosphatase (PHP family) | 1.45E-17 | 5.1E-16 | 72.8 | 0.368 | 0.724 | 0.357 |
| K03738 | aldehyde:ferredoxin oxidoreductase | 1.73E-17 | 6.02E-16 | 72.4 | 0.282 | 1.04 | 0.753 |
| K03856 | 3-deoxy-7-phosphoheptulonate synthase | 2.1E-17 | 7.15E-16 | 72 | 0.322 | 0.713 | 0.392 |
| K02025 | multiple sugar transport system permease protein | 7.46E-16 | 2.03E-14 | 65 | 0.641 | 2.19 | 1.55 |
| K02026 | multiple sugar transport system permease protein | 3.12E-14 | 6.91E-13 | 57.7 | 0.672 | 2.19 | 1.52 |

| | | | | | | | |
|---|---|---|---|---|---|---|---|
| K01443 | N-acetylglucosamine-6-phosphate deacetylase | 5.1E-14 | 1.08E-12 | 56.7 | 0.337 | 0.867 | 0.53 |
| K07030 | uncharacterized protein | 1.21E-12 | 2.07E-11 | 50.5 | 0.972 | 2.17 | 1.2 |
| K07814 | putative two-component system response regulator | 5.02E-11 | 6.75E-10 | 43.2 | 0.753 | 1.28 | 0.523 |
| K02027 | multiple sugar transport system substrate-binding protein | 6.44E-11 | 8.55E-10 | 42.7 | 0.562 | 2.26 | 1.7 |

**Supplementary Table 2. Significant KOs among partners vs. non-partners.** Summary of KOs found to be significantly overrepresented in those dMAGs identified as partners of methanogens (partners), over dMAGs that do not correlate with methanogens (non-partners) defined as $r < 0.5$ with any methanogen across all reactors. The p-values (p) were determined from a non-parametric Kruskal-Wallis test (statistic tau) comparing individual KO frequencies in the two groups and Benjamini-Hochberg adjusted p-values calculated to account for multiple comparisons (q). Diff. is the difference (Mean partner – Mean non-partner) between the mean gene frequencies in the two groups.

| | FALSE | TRUE | Class error |
|---|---|---|---|
| **FALSE** | 352 | 105 | 0.2297 |
| **TRUE** | 45 | 405 | 0.1000 |

**Supplementary Table 3. Random forest results relating to partner vs. non-partner distinction.** Outcome statistics, i.e. confusion matrix of a random forest based classification of all bacterial dMAGs into partner vs. non-partner categories based on their KO content. There were 500 trees built, with 49 variables tried at each split. Estimated error rate was 16.54%, i.e. 83.46% accuracy.

| KO id | KO name | p | q | tau | Mean non-partners | Mean partners | Diff. |
|---|---|---|---|---|---|---|---|
| K07074 | Uncharactherised protein | 2.63e-07 | 0.000389 | 26.5 | 0.0 | 0.352 | 0.353 |
| K02834 | Ribosome-binding factor A | 1.93e-05 | 0.0143 | 18.3 | 0.245 | 0.764 | 0.519 |
| K09797 | Uncharactherised protein | 5.94e-05 | 0.0294 | 16.1 | 0.072 | 0.411 | 0.339 |
| K00970 | Poly(A) polymerase | 1.23e-04 | 0.0451 | 14.7 | 0.036 | 0.294 | 0.258 |
| K02939 | Large subunit ribosomal protein L9 | 2.75e-04 | 0.0451 | 13.2 | 0.272 | 0.705 | 0.433 |
| K02117 | V/A type H+/Na+ transporting ATPase subunit A | 3.04e-04 | 0.0451 | 13.0 | 0.0 | 0.117 | 0.118 |
| K02437 | Glycine cleavage system H protein | 3.04e-04 | 0.0451 | 13.0 | 0.0 | 0.117 | 0.118 |
| K03547 | DNA repair protein SbcD/Mre11 | 3.04e-04 | 0.0451 | 13.0 | 0.0 | 0.117 | 0.118 |
| K07029 | Diacylglycerol kinase (ATP) | 3.04e-04 | 0.0451 | 13.0 | 0.0 | 0.117 | 0.118 |
| K07494 | Putative transposase | 3.04e-04 | 0.0451 | 13.0 | 0.0 | 0.117 | 0.118 |

| | | | | | | | |
|---|---|---|---|---|---|---|---|
| K03282 | Large conductance mechanosensitive channel | 4.3e-04 | 0.058 | 12.4 | 0.755 | 0.176 | -0.569 |

**Supplementary Table 4. Significant KOs in Patescibacteria dMAGs that are predicted partners of methanogens.** KOs and their associated statistics test results for being enriched in Patescibacteria that are predicted to associate with methanogens compared to those that do not. The p-values (p) were determined from a non-parametric Kruskal-Wallis test (test statistic tau) comparing individual KO frequencies in the two groups and Benjamini-Hochberg adjusted p-values calculated to account for multiple comparisons (q). Diff. is the difference (Mean partner – Mean non-partner) between the mean gene frequencies in the two groups.

| **AD id** | **nSamples** | **R2** | **p** |
|---|---|---|---|
| AD2.1 | 33 | 0.355 | 0.001 |
| AD2.2 | 26 | 0.493 | 0.001 |
| AD3 | 14 | 0.535 | 0.001 |
| AD7 | 44 | 0.418 | 0.001 |
| AD10 | 36 | 0.184 | 0.002 |
| AD11 | 35 | 0.374 | 0.001 |
| AD12 | 34 | 0.343 | 0.001 |
| AD16.1 | 26 | 0.329 | 0.001 |
| AD16.2 | 30 | 0.258 | 0.001 |
| AD18D | 21 | 0.459 | 0.001 |
| AD18H | 19 | 0.281 | 0.001 |
| AD20 | 32 | 0.128 | 0.011 |

**Supplementary Table 5. ANOVA of community structure against sampling week.** A multivariate permutation ANOVA (as implemented in the adonis function of vegan) of community structure (measured by normalised dMAG coverage) against sampling week was performed for each reactor independently using Bray-Curtis distances. We give the proportion of variance explained (R2) and p-value (p).

| AD id | Variable | R2 | p |
|---|---|---|---|
| AD10 | week | 0.22 | 0.001 |
| AD10 | o2... | 0.053 | 0.013 |
| AD10 | temperature | 0.14 | 0.001 |
| AD10 | beet.T. | 0.089 | 0.001 |
| AD10 | beet_pulp.T. | 0.084 | 0.001 |
| AD10 | pH | 0.095 | 0.002 |
| AD11 | week | 0.35 | 0.001 |
| AD11 | beet.T. | 0.08 | 0.001 |
| AD11 | beet_pulp.T. | 0.17 | 0.001 |
| AD11 | chicken_manure.T. | 0.044 | 0.005 |
| AD11 | grass_silage.T. | 0.028 | 0.038 |
| AD11 | pH | 0.03 | 0.022 |
| AD12 | week | 0.34 | 0.001 |
| AD12 | beet_pulp.T. | 0.19 | 0.001 |
| AD12 | grass_silage.T. | 0.03 | 0.034 |
| AD12 | maize_silage.T. | 0.065 | 0.002 |
| AD12 | pig_manure.T. | 0.078 | 0.001 |
| AD16.2 | week | 0.2 | 0.003 |
| AD16.2 | TotalF | 0.17 | 0.004 |
| AD18D | week | 0.48 | 0.001 |
| AD18H | week | 0.29 | 0.001 |
| AD18H | temperature | 0.19 | 0.001 |
| AD18H | beet.T. | 0.064 | 0.004 |
| AD18H | chicken_manure.T. | 0.14 | 0.001 |
| AD18H | grass_silage.T. | 0.058 | 0.022 |
| AD2.1 | week | 0.35 | 0.001 |
| AD2.1 | temperature | 0.17 | 0.001 |
| AD2.2 | week | 0.49 | 0.001 |
| AD2.2 | temperature | 0.067 | 0.01 |
| AD2.2 | maize_silage.T. | 0.06 | 0.021 |
| AD7 | week | 0.42 | 0.001 |
| AD7 | o2... | 0.095 | 0.001 |
| AD7 | whey.T. | 0.035 | 0.013 |
| AD7 | TotalF | 0.037 | 0.016 |

**Supplementary Table 6. Multivariate ANOVA of community structure against different metadata.** A multivariate permutation ANOVA (as implemented in the adonis function of vegan) of community structure (measured by normalised dMAG coverage) against sampling week and all additional metadata variables that were at least 90% complete for that reactor, was performed for each reactor independently using Bray-Curtis distances. We give the proportion of variance explained (R2), p-values (p) but only for those variables with $p < 0.05$.

| AD id | nSamples | R2 | p |
|---|---|---|---|
| AD2.1 | 33 | 0.173 | 0.012 |
| AD2.2 | 26 | 0.0599 | 0.219 |
| AD3 | 14 | 0.107 | 0.242 |
| AD7 | 44 | 0.103 | 0.013 |
| AD10 | 36 | 0.0434 | 0.209 |
| AD11 | 35 | 0.382 | 0.001 |
| AD12 | 34 | 0.311 | 0.001 |
| AD16.1 | 26 | 0.421 | 0.001 |
| AD16.2 | 30 | 0.0545 | 0.187 |
| AD18D | 21 | 0.506 | 0.001 |
| AD18H | 19 | 0.651 | 0.001 |
| AD20 | 32 | 0.0826 | 0.093 |

**Supplementary Table 7. Multivariate ANOVA of methanogenic functional trait abundance against sampling week.** A multivariate permutation ANOVA (as implemented in the adonis function of vegan) of methanogenesis trait abundances (measured by aggregate normalised dMAG coverage) against sampling week was performed for each reactor independently using Bray-Curtis distances. We give the proportion of variance explained (R2), p-values (p).

| Variable | Estimate | Std. Error | t value | Pr(>\|t\|) |
|---|---|---|---|---|
| (Intercept) | 530.008 | 203.461 | 2.605 | 0.00995 ** |
| AD12 | 14.611 | 16.389 | 0.891 | 0.37385 |
| AD16.2 | 420.771 | 18.738 | 22.456 | < 2e-16 *** |
| AD18D | 136.053 | 65.603 | 2.074 | 0.03950 * |
| AD2.1 | 373.026 | 152 | 26.358 | < 2e-16 *** |
| AD2.2 | 347.245 | 15.511 | 22.387 | < 2e-16 *** |
| ADAD7 | 313.702 | 14.425 | 21.747 | < 2e-16 *** |
| temperature | 15.297 | 5.032 | 3.040 | 0.00272 ** |

**Supplementary Table 8. Multivariate regression of yield against temperature and AD reactor identity.** Yield was predicted as a function of temperature with AD reactor as an additional discrete variable. Multiple R-squared: 0.8843, Adjusted R-squared: 0.8798 F-statistic: 197.6 on 7 and 181 DF, p-value: < 2.2e-16

| MAG | chi | p | q | Class | Genus | Partner |
|---|---|---|---|---|---|---|
| AD7_Bin_1857 | 98.3 | 1.69E-16 | 9.93E-14 | / | / | TRUE |
| AD3_Bin_901 | -104 | 3.96E-16 | 1.17E-13 | Clostridia | DTU059 | FALSE |
| AD12_Bin_716 | 69.9 | 8.7E-16 | 1.7E-13 | Dethiobacteria | UBA8154 | TRUE |
| AD7_Bin_553 | 44.9 | 4.35E-13 | 5.16E-11 | Cloacimonadia | Cloacimonas | TRUE |
| AD7_Bin_565 | -70.5 | 4.39E-13 | 5.16E-11 | Paceibacteria | / | FALSE |
| AD7_Bin_494 | 108 | 7.18E-13 | 7.04E-11 | Thermotogae | Mesotoga | TRUE |
| AD7_Bin_1785 | -42.1 | 9.38E-12 | 7.88E-10 | Bacilli | / | TRUE |
| MINI_Bin_19 | 39.9 | 1.43E-11 | 1.05E-09 | JS1 | / | TRUE |
| AD3_Bin_291 | 65 | 2.87E-11 | 1.87E-09 | Syntrophia | UBA4810 | FALSE |
| AD2.1_Bin_1236 | 51.3 | 3.75E-11 | 2.16E-09 | Desulfomonilia | UBA1062 | FALSE |
| AD2.1_Bin_616 | 86.2 | 4.05E-11 | 2.16E-09 | Caldisericia | / | FALSE |
| AD2.2_Bin_1085 | 74.9 | 4.56E-11 | 2.21E-09 | Thermococci | Methanofastidiosum | FALSE |
| AD2.1_Bin_973 | -42.6 | 4.9E-11 | 2.21E-09 | Spirochaetia | UBA9732 | FALSE |
| AD7_Bin_568 | 85.6 | 8.79E-11 | 3.69E-09 | UBA5377 | UBA1398 | TRUE |
| AD2.1_Bin_491 | -47.2 | 1.28E-10 | 5.03E-09 | Syntrophomonadia | Syntrophaceticus | TRUE |
| AD10_Bin_1032 | 96.8 | 2.71E-10 | 9.97E-09 | Microgenomatia | UBA6130 | FALSE |
| AD7_Bin_750 | 60.3 | 4.99E-10 | 1.73E-08 | Bacilli | DTU067 | FALSE |
| AD18H_Bin_461 | 102 | 7.01E-10 | 2.29E-08 | Clostridia | Herbivorax | FALSE |
| AD2.2_Bin_692 | 89.8 | 1.3E-09 | 4.02E-08 | Dethiobacteria | UBA9862 | FALSE |

| | | | | | | |
|---|---|---|---|---|---|---|
| AD2.2_Bin_495 | -34.3 | 1.43E-09 | 4.21E-08 | Bacteroidia | F082 | TRUE |

**Supplementary Table 9. Top 20 dMAGs showing significant correlations with yield (i.e. reactor performance).** Results (**chi**) are correlation coefficients from multivariate regressions of reactor yield (methane volume produced per tonne of feed) against log dMAG normalised abundance together with reactor identity and operating temperature. The p-values (p) and adjusted p-values (q) for the dMAG coefficient are also given together with the class and genus of the dMAG and whether it is a potential methanogenic partner as per our definition ($r > 0.9$ and $q < 1.0e-5$).

| | Not positively strongly associated with yield | Positively strongly associated with yield | |
|---|---|---|---|
| Partner - False | 442 | 65 | 65/507 = 12.8% |
| Partner - True | 60 | 21 | 21/81 = 25.9 % |
| | 60/502 = 11.9% | 21/86 = 24.4% | |

**Supplementary Table 10. Confusion matrix between dMAGs that correlate significantly ($q < 0.001$) and positively (chi > 0) with yield and those that are classified as methanogenic partners.** The association was significant with p-value = 0.003 (Fisher's exact test).

**SUPPLEMENTARY FIGURES**

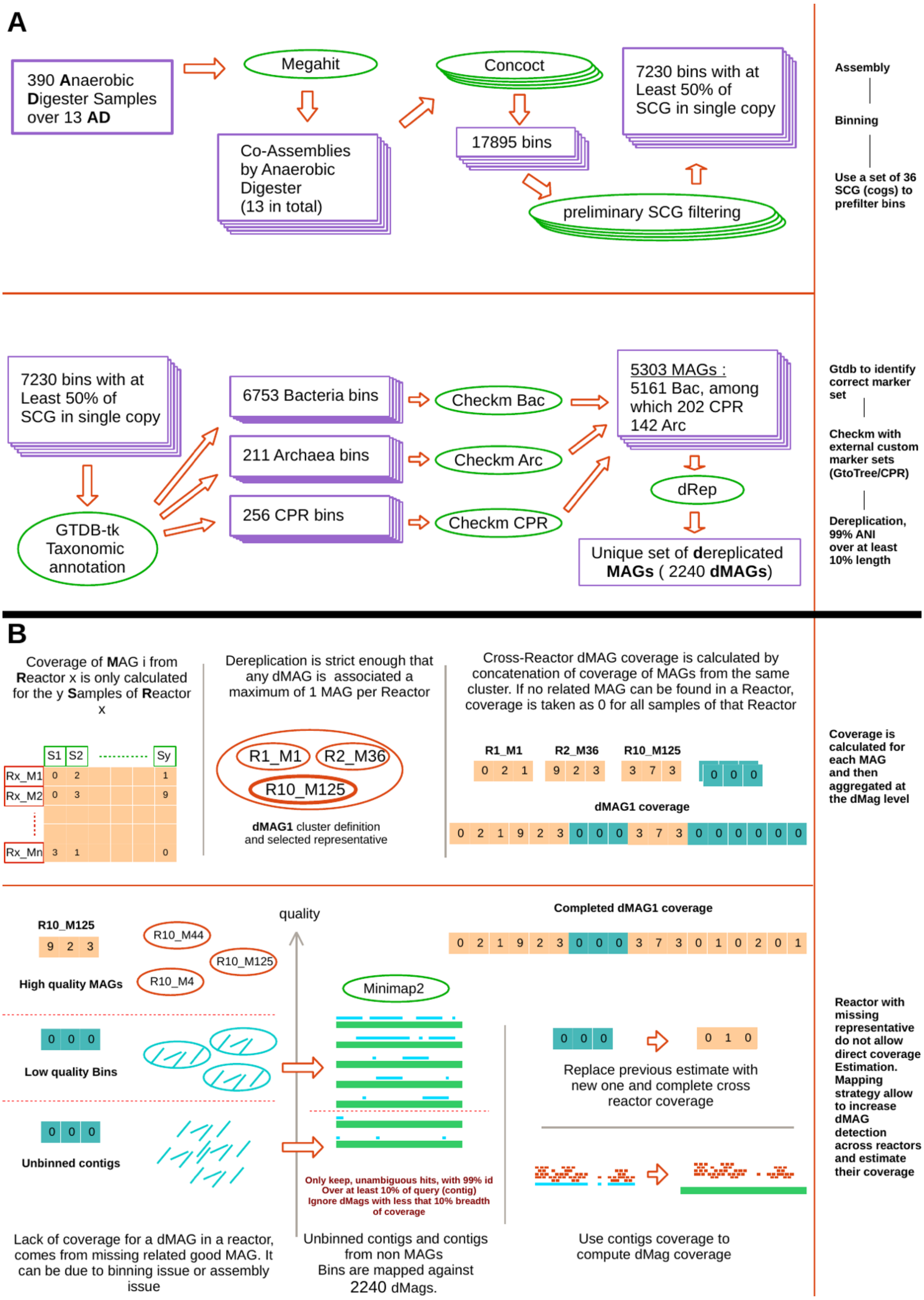


**Supplementary Figure 1. Building a MAG collections and coverage profile across reactors. (A)** Bins are built using concoct, quality is assessed through appropriate marker sets to select MAGs and MAGs dereplicated into dMAGs using dRep. **(B)** dMAG coverage is defined by the coverage of the corresponding MAGs in the dRep cluster and additionally through mapping of contigs to dMAGs with minimap2.

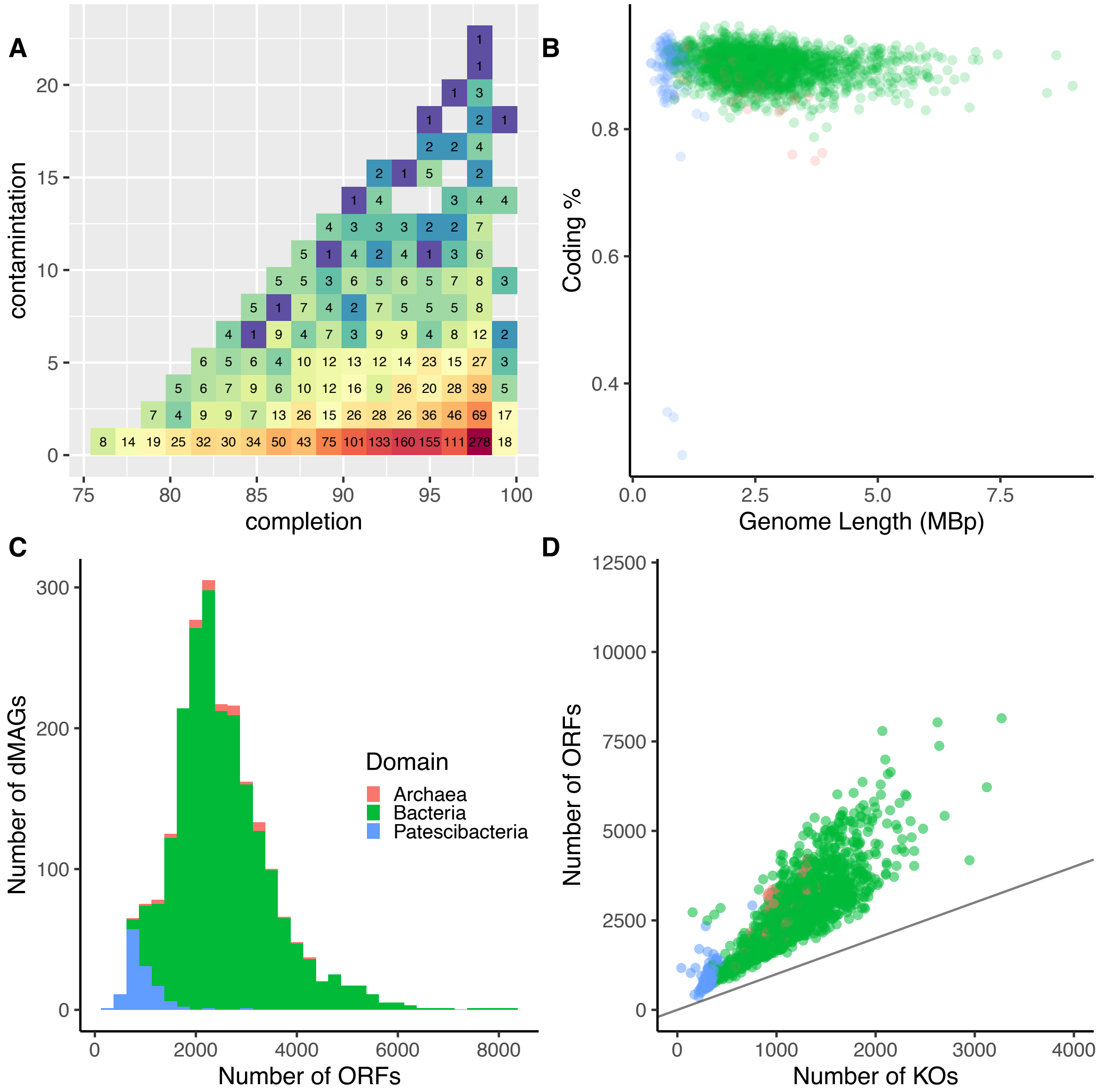


**Supplementary Figure 2. Statistics for dMAG properties.** General information on dMAGs identified in this study. **(A)** Contamination and completion of all dMAGs. **(B)** Genome length of dMAGs against their coding percentage. **(C)** Distribution of dMAGs according to the number of open reading frames (ORFs) that they contain. **(D)** Analysis of number of KEGG orthologs (KOs) against ORFs in each dMAG. For panels B to D, Archaea, bacteria and Patescibacteria (i.e. CPR) are colored in red, green, and blue respectively.

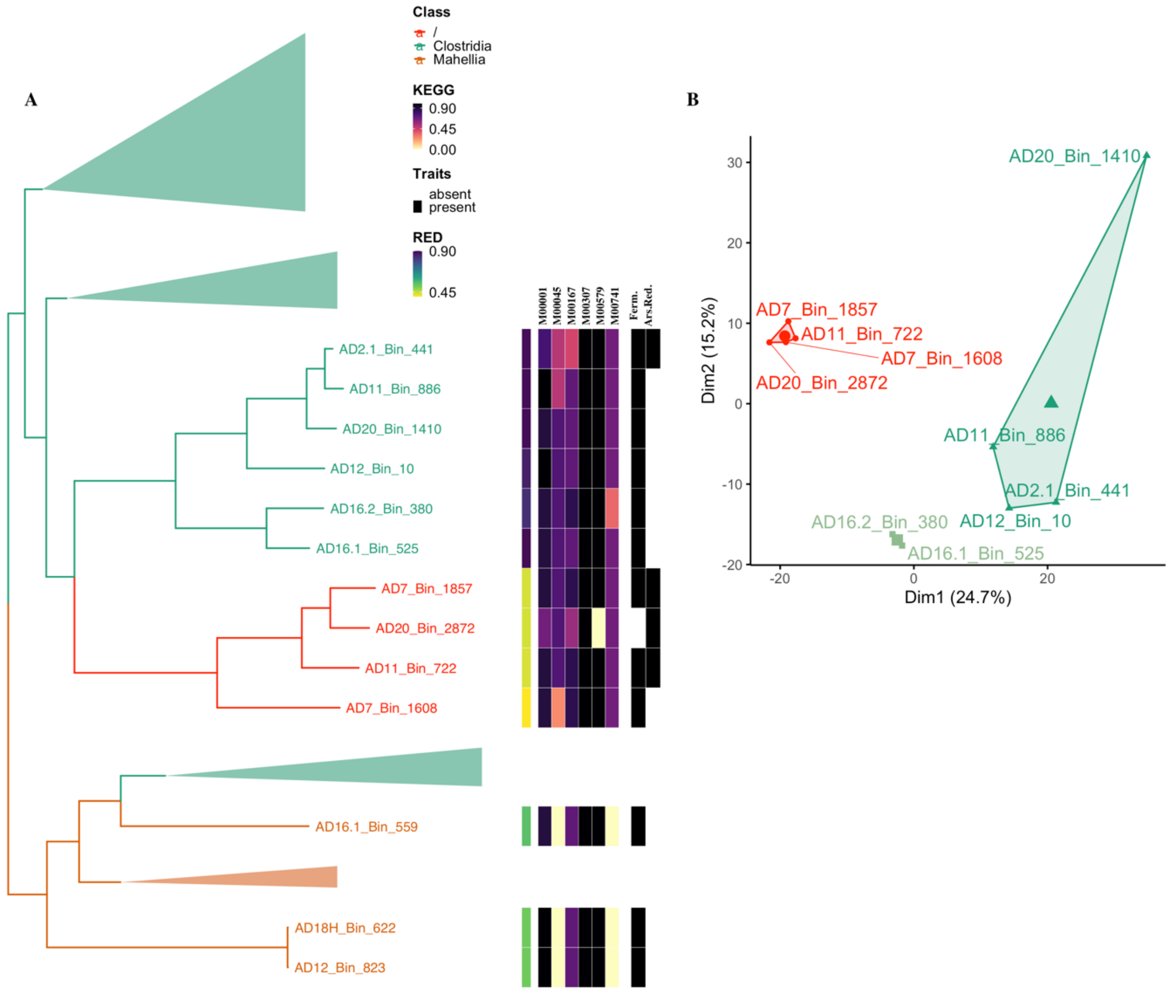


**Supplementary Figure 3. Phylogenetic tree for novel class with four dMAGs within Firmicutes phylum. (A)** Sub-clade of the *de novo* phylogenetic tree constructed from all bacterial dMAGs and family representatives from the GTDB (i.e. part of the tree shown in Fig. 1). Branches are coloured according to GTDB class assignment. Alongside the tree, we show the RED value, KEGG module completeness, and fermentation and arsenate reduction trait assignment for each dMAG. The KEGG module completeness (see *Methods*) is shown for the modules: glycolysis (M00001), pyruvate-to-acetylCoA conversion (M00307), reductive pentose phosphate cycle (M00167), and acetate production (M00579) pathways, histidine degradation (M00045), and propanoyl-CoA pathways (M00741). **(B)** Results of a clustering analysis using all annotated KEGG orthologs (KOs) of each dMAG (see *Methods*).

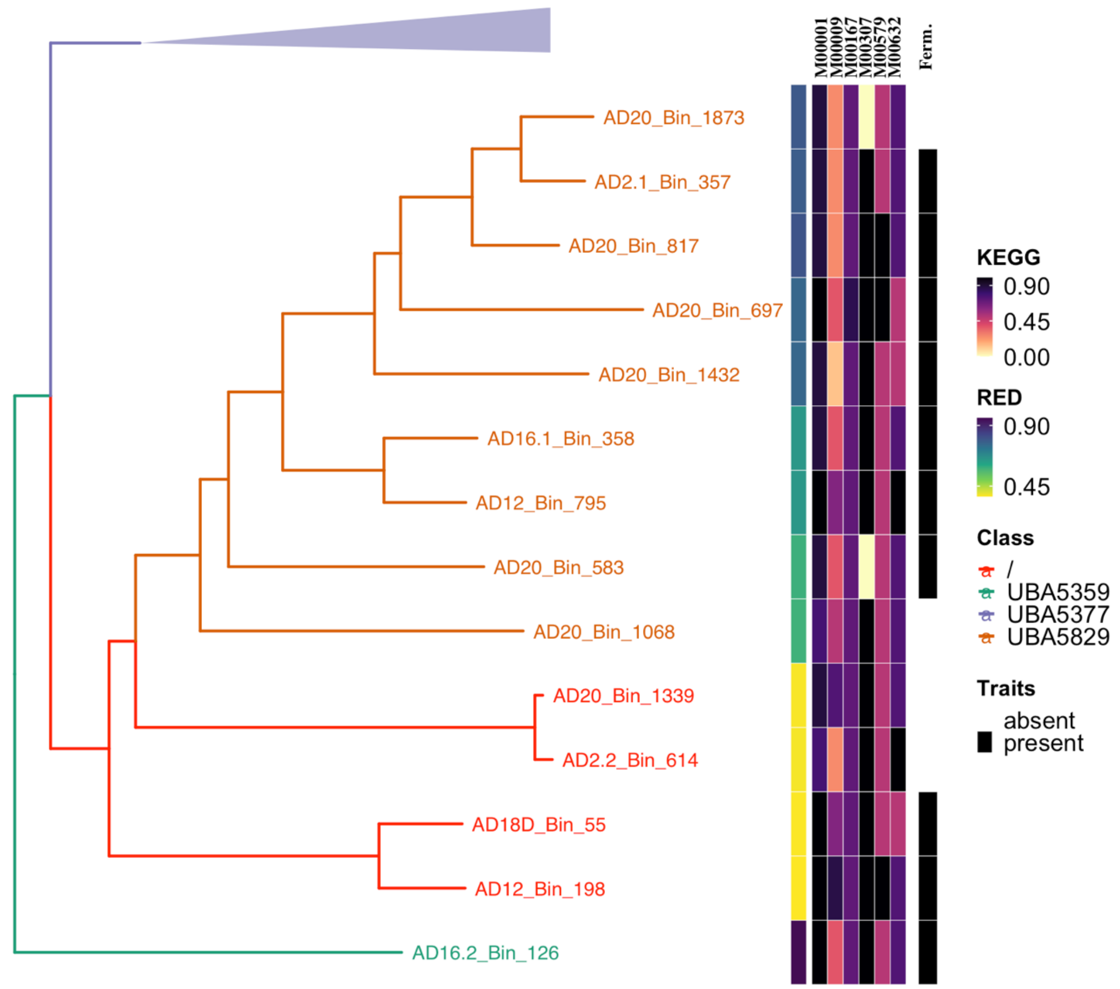


**Supplementary Figure 4. Phylogenetic tree for novel class with four dMAGs within Armatimonadota phylum.** Sub-clade of the *de novo* tree constructed from all bacterial dMAGs and family representatives from the GTDB (i.e. part of the tree shown in Fig. 1). Branches are coloured according to GTDB class assignment. Alongside the tree, we show the RED value, KEGG module completeness, and fermentation trait assignment for each dMAG. The KEGG module completeness (see *Methods*) is shown for the modules: glycolysis (M00001), pyruvate-to-acetylCoA conversion (M00307), reductive pentose phosphate cycle (M00167), and acetate production (M00579) pathways, the tricarboxylic acid (TCA) module (M00009), and galactose degradation (M00632).

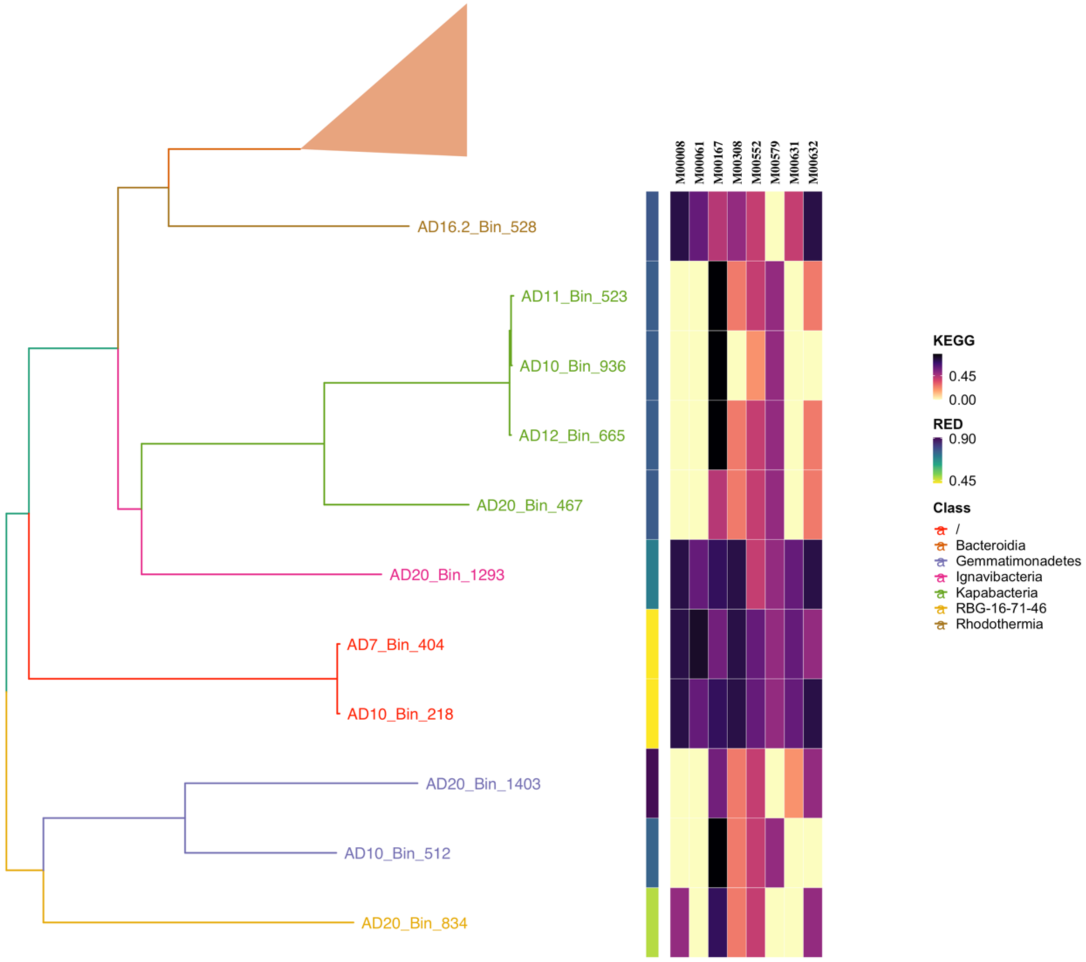


**Supplementary Figure 5. Phylogenetic tree for novel class with four dMAGs within Latescibacterota phylum.** Sub-clade of the *de novo* tree constructed from all bacterial dMAGs and family representatives from the GTDB (i.e. part of the tree shown in Fig. 1). Branches are coloured according to GTDB class assignment. Alongside the tree, we show the RED value and KEGG module completeness. The KEGG module completeness (see *Methods*) is shown for the modules: Entner-Doudoroff pathway (M00008), reductive pentose phosphate cycle (M00167), gluconate to glycerate-3P conversion (M00308), and acetate production (M00579) pathways, and D-Gluocuronate (M00061), D-Galactonate (M00552), D-Galacturonate (M00631), and Galactose (M00632) degradation.

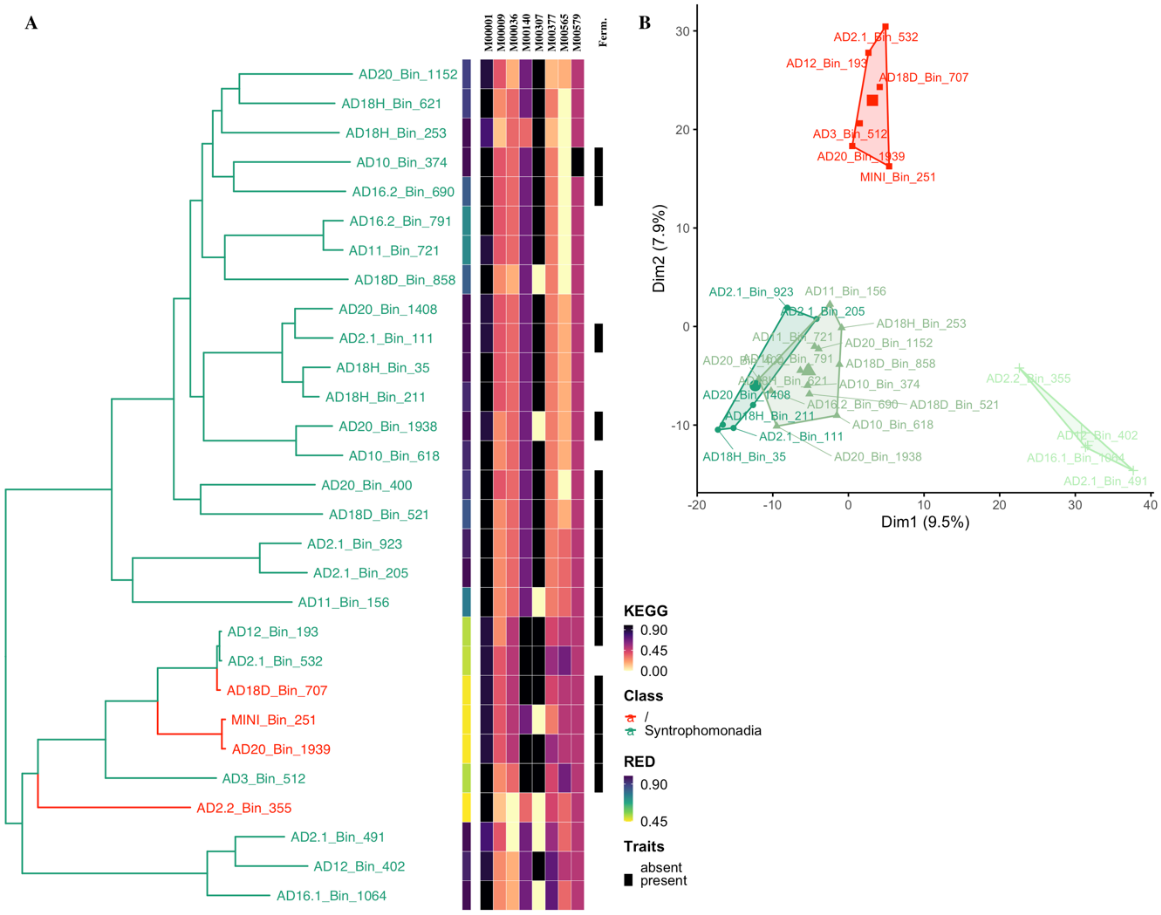

**Supplementary Figure 6. Phylogenetic tree for novel class with four dMAGs within Syntrophomonadia class. (A)** Sub-clade of the *de novo* tree constructed from all bacterial dMAGs and family representatives from the GTDB (i.e. part of the tree shown in Fig. 1). Branches are colored according to GTDB class assignment. Alongside the tree, we show the RED value, KEGG module completeness, and fermentation trait assignment for each dMAG. The KEGG module completeness (see *Methods*) is shown for the modules: glycolysis (M00001), pyruvate-to-acetylCoA conversion (M00307), reductive acetylCoA (Wood-Ljungdahl) (M00377), and acetate production (M00579) pathways, the tricarboxylic acid (TCA) module (M00009), leucine degradation (M00036), trehalose biosynthesis (M00565) and C1 conversion pathways (M00140). **(B)** Results of a clustering analysis using all annotated KEGG orthologs (KOs) of each dMAG (see *Methods*).

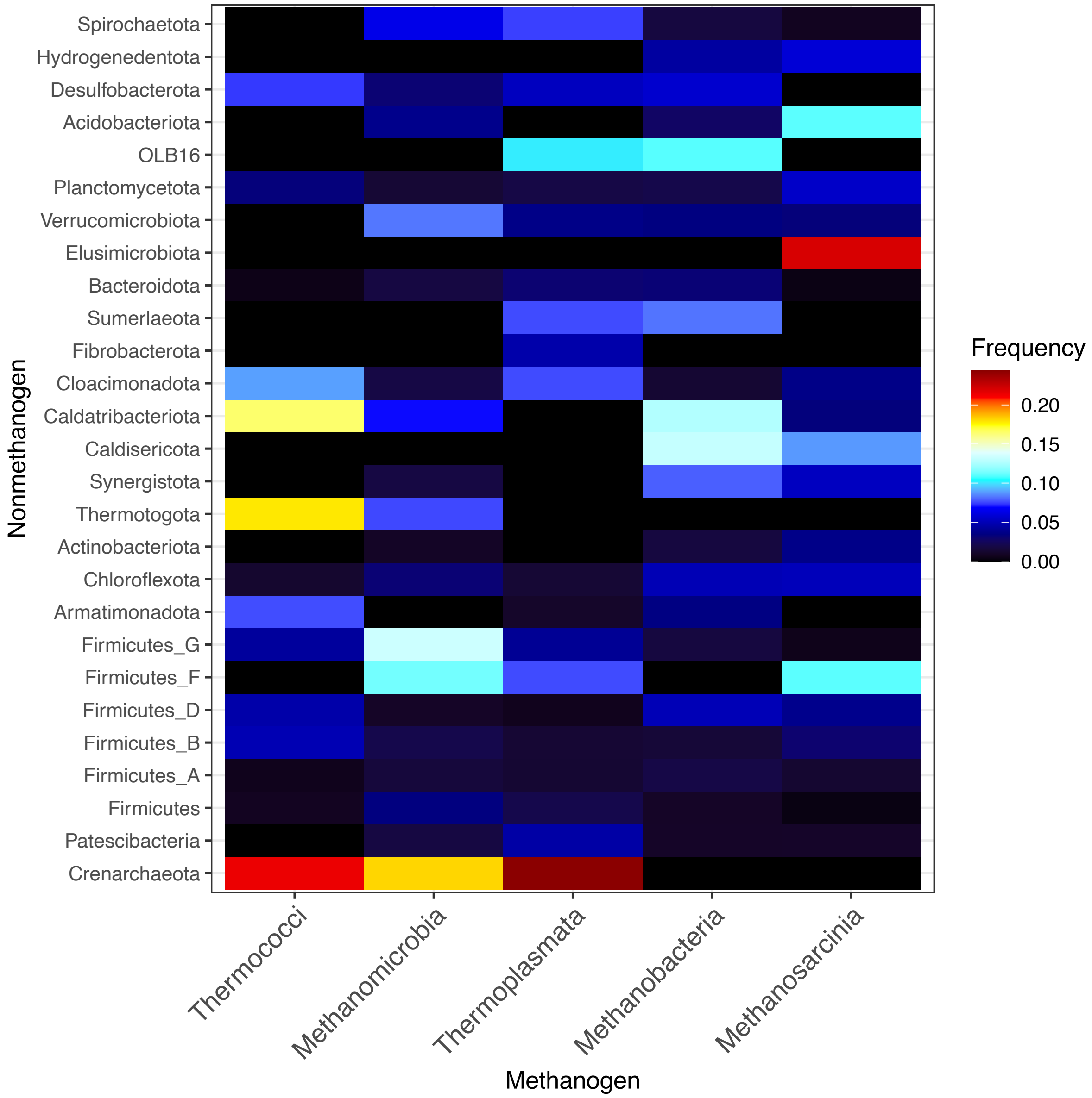


**Supplementary Figure 7. Distribution of predicted syntrophs among taxa.** The frequency with which dMAGs in non-methanogenic phyla interact with different methanogen classes. Frequencies are defined by dividing the total number of interactions between a phylum and a methanogen class by the number of dMAGs in the phylum to normalise for differences in dMAG richness. These frequencies are then normalised to sum to one in each column i.e. across a methanogen class. The association between phyla and class was highly significant (p-value = 6.829e-10, Chi-squared test).

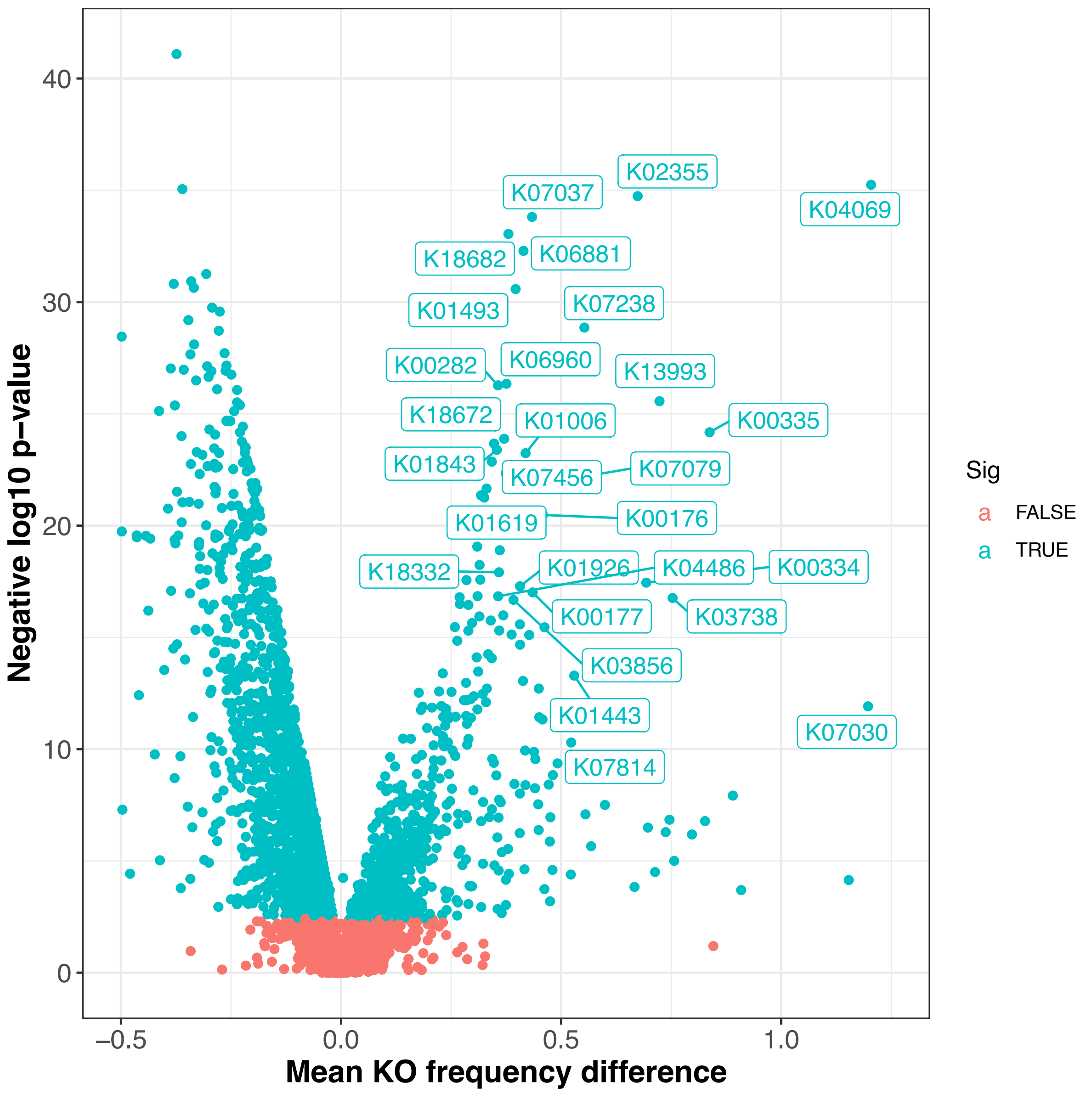


**Supplementary Figure 8. Volcano plot for KEGG orthologs (KOs) that are significantly overrepresented in predicted syntrophs (methanogen partners) against non-partners.** Each point on the graph shows a specific KO, with the x-axis the frequency in partners minus non-partners and the y-axis the negative log10 of the p-value in a Kruskal-Wallis test comparing individual KO frequencies in the two groups. Significant KOs with a Benjamini-Hochberg adjusted p-value < 0.05 are highlighted. KO names are given in Supplementary Table S2.

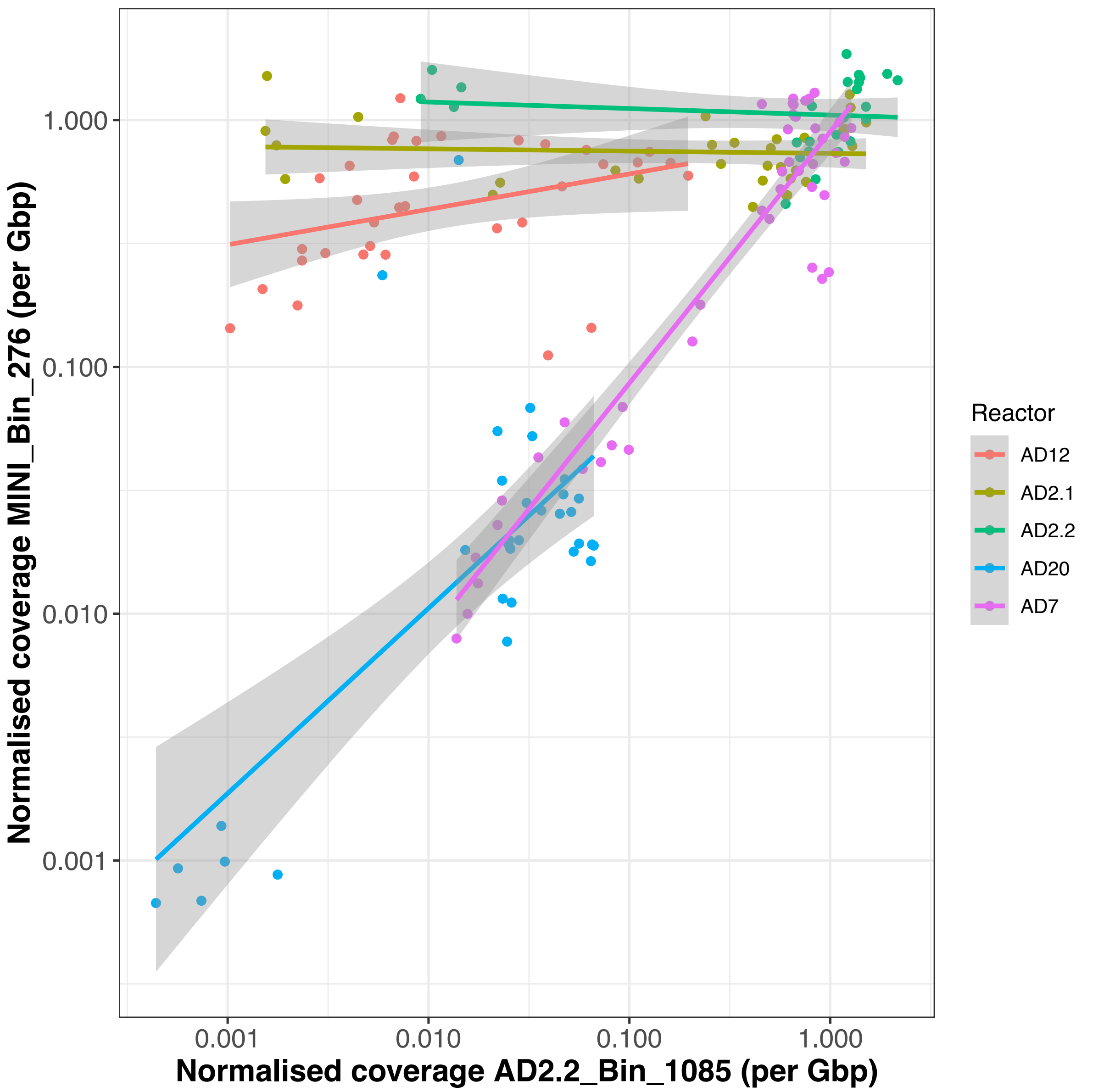


**Supplementary Figure 9. Coverage correlations across all reactors they cooccur in for a predicted syntrophic pair.** The methanogenic Thermococci (AD2.2_Bin_1085, *Methanofastidiosum*), and the Bathyarchaeia (MINI_Bin_276) are predicted to interact based on a strong correlation in AD7 ($r = 0.95$, p-value = 4.59e-22). Here we show there normalised coverages across all five reactors they cooccur in. In AD20 they may also be associated ($r = 0.72$, p-value = 3.2e-6) but in the three others (AD2.1, AD2.2 and AD12) they have no significant association ($p > 0.1$).

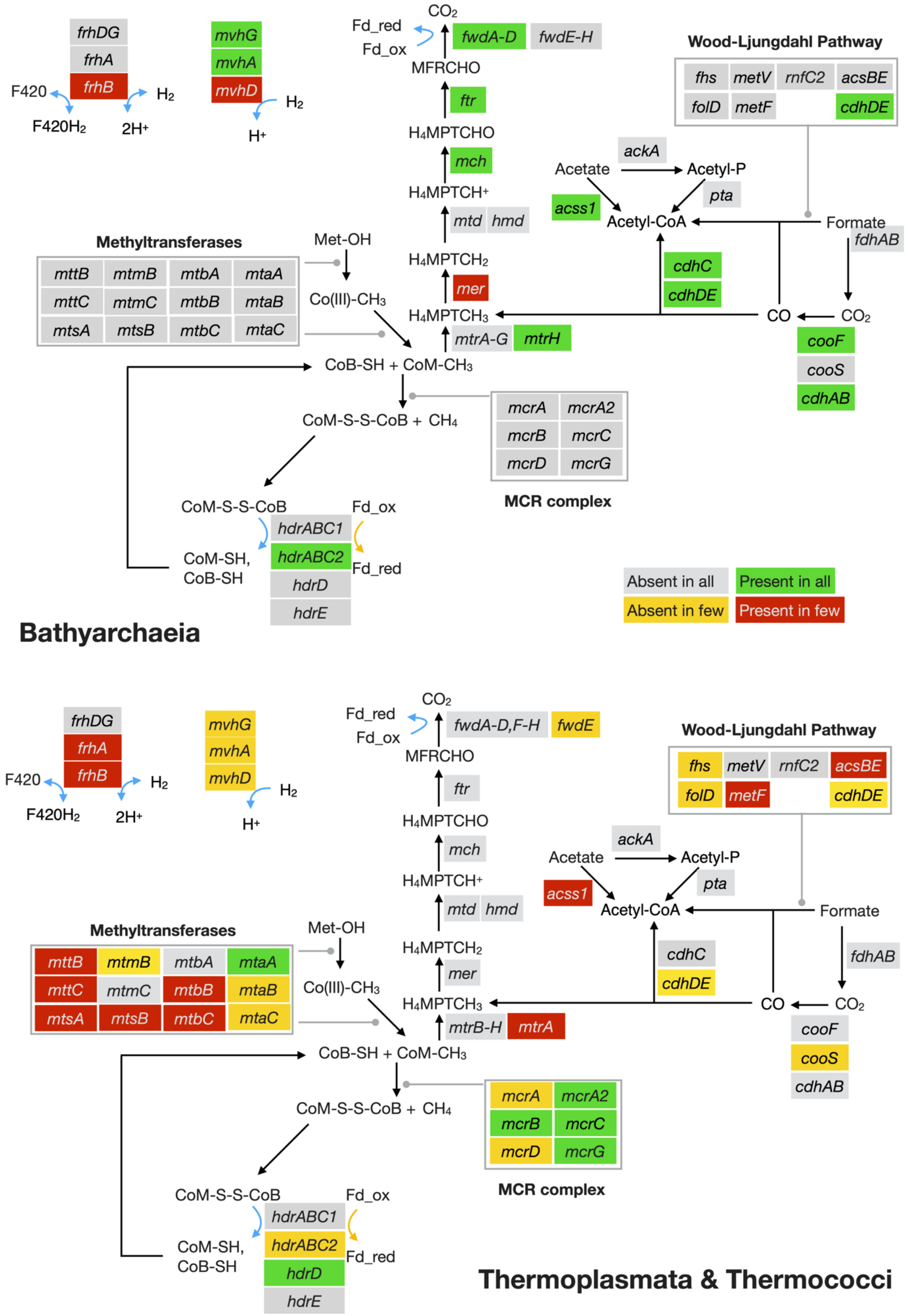


**Supplementary Figure 10. Metabolic mapping of select KOs in Bathyarchaeia and Thermoplasmata and Thermococci dMAGs.** The cartoon figure shows the presence or absence of key genes involved in methylotrophic methanogenesis and reductive acetate (i.e. Wood-Ljungdahl) pathways – see also *Supplementary File 4*. The analysis is based on the following dMAGs; AD2.2_Bin_1085 (Thermococci); AD11_Bin_772, MINI_Bin_140, AD3_Bin_264, AD20_Bin_1426, AD7_Bin_314, AD7_Bin_584 (Thermoplasmata); and MINI_Bin_276, AD2.2_Bin624, AD7_Bin_974, AD7_Bin_127 (Bathyarchaeia).

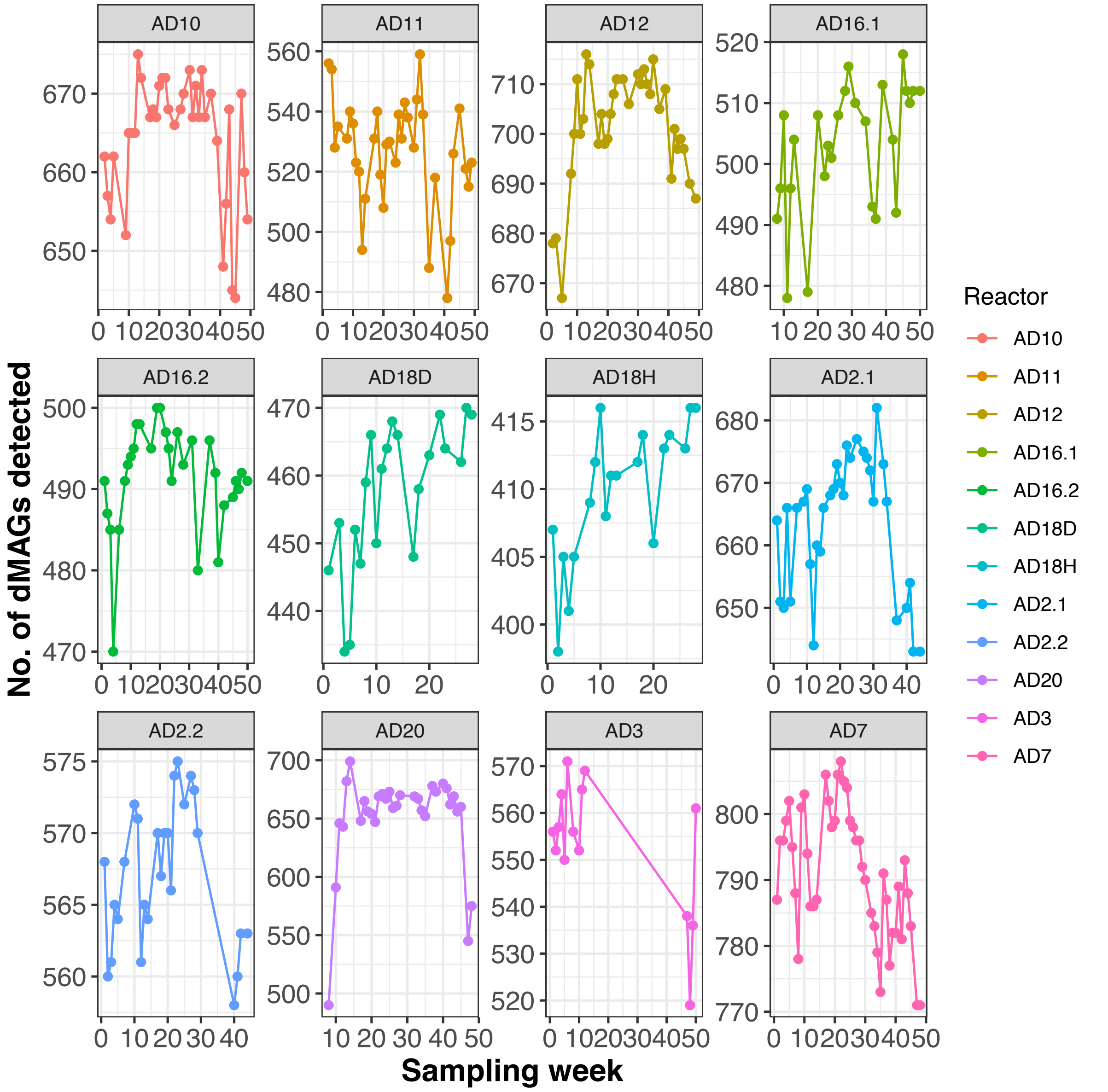


**Supplementary Figure 11. Total dMAG richness for each reactor over time.** The number of dMAGs with non-zero coverage detected at each sampling time.

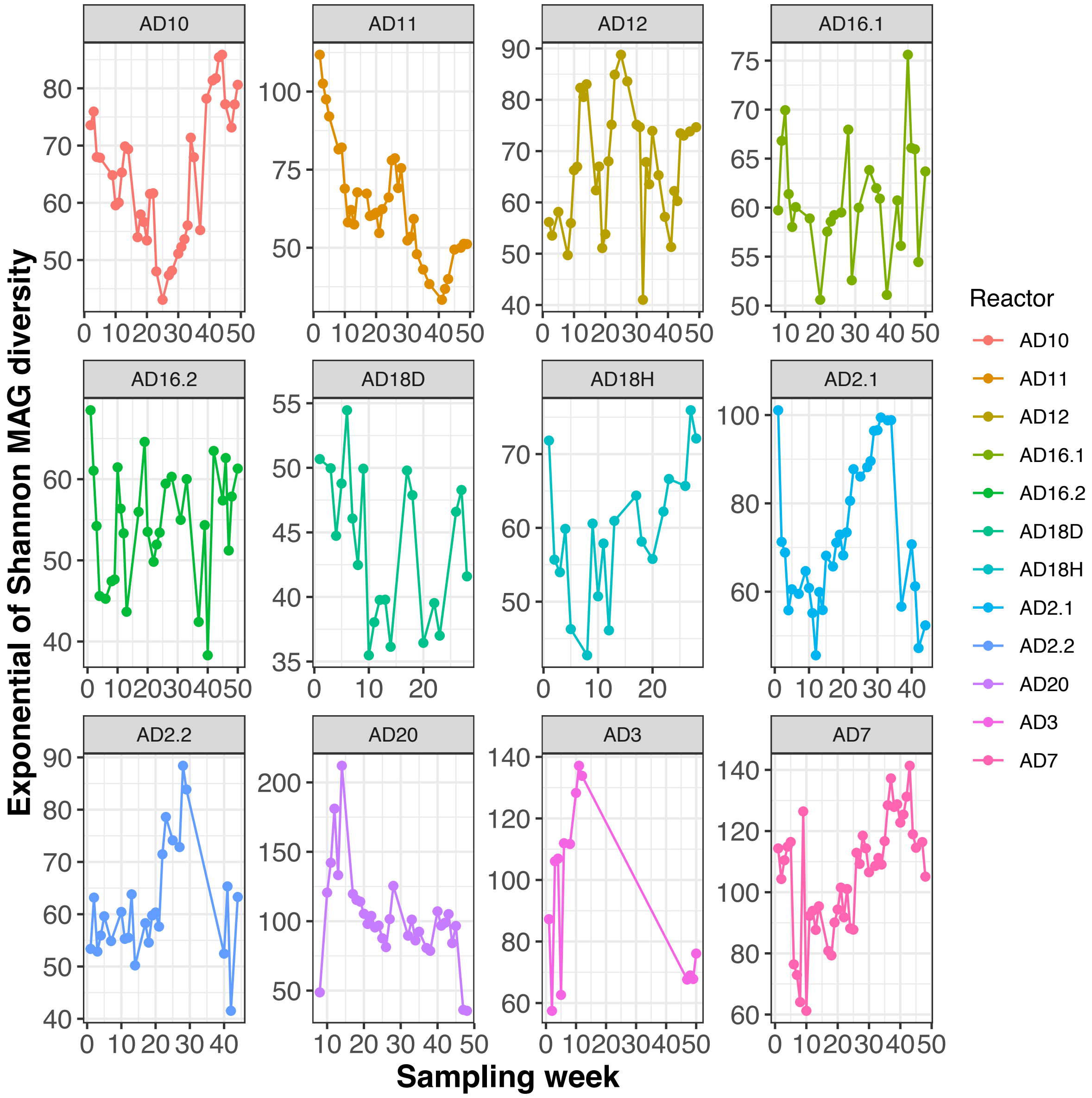


**Supplementary Figure 12. Shannon diversity of dMAGs distribution for each reactor over time.** The Shannon diversity of normalised dMAG coverage profile was calculated at each sample time and raised to the exponential. Shannon diversity decreased significantly over time in AD11 ($p = 2.1e-08$) and AD20 ($p = 1.0e-3$) but increased in AD7 ($p = 4.8e-05$).

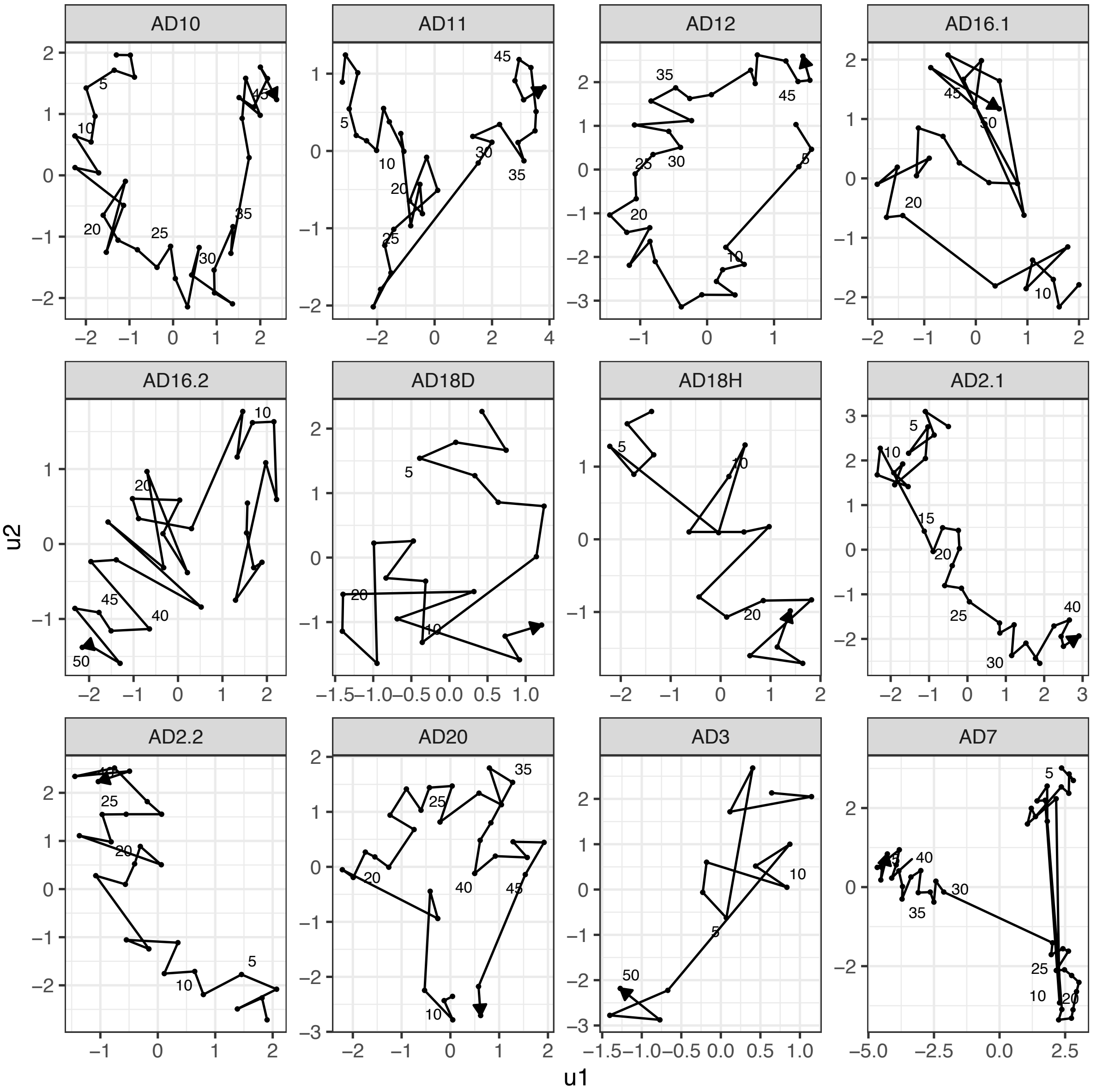


**Supplementary Figure 13. Changing community dMAG structure for each reactor over time.** Normalised dMAG coverage profiles (2240 dimensions) were log-transformed and then mapped to two dimensions using the umap dimension reduction algorithm.

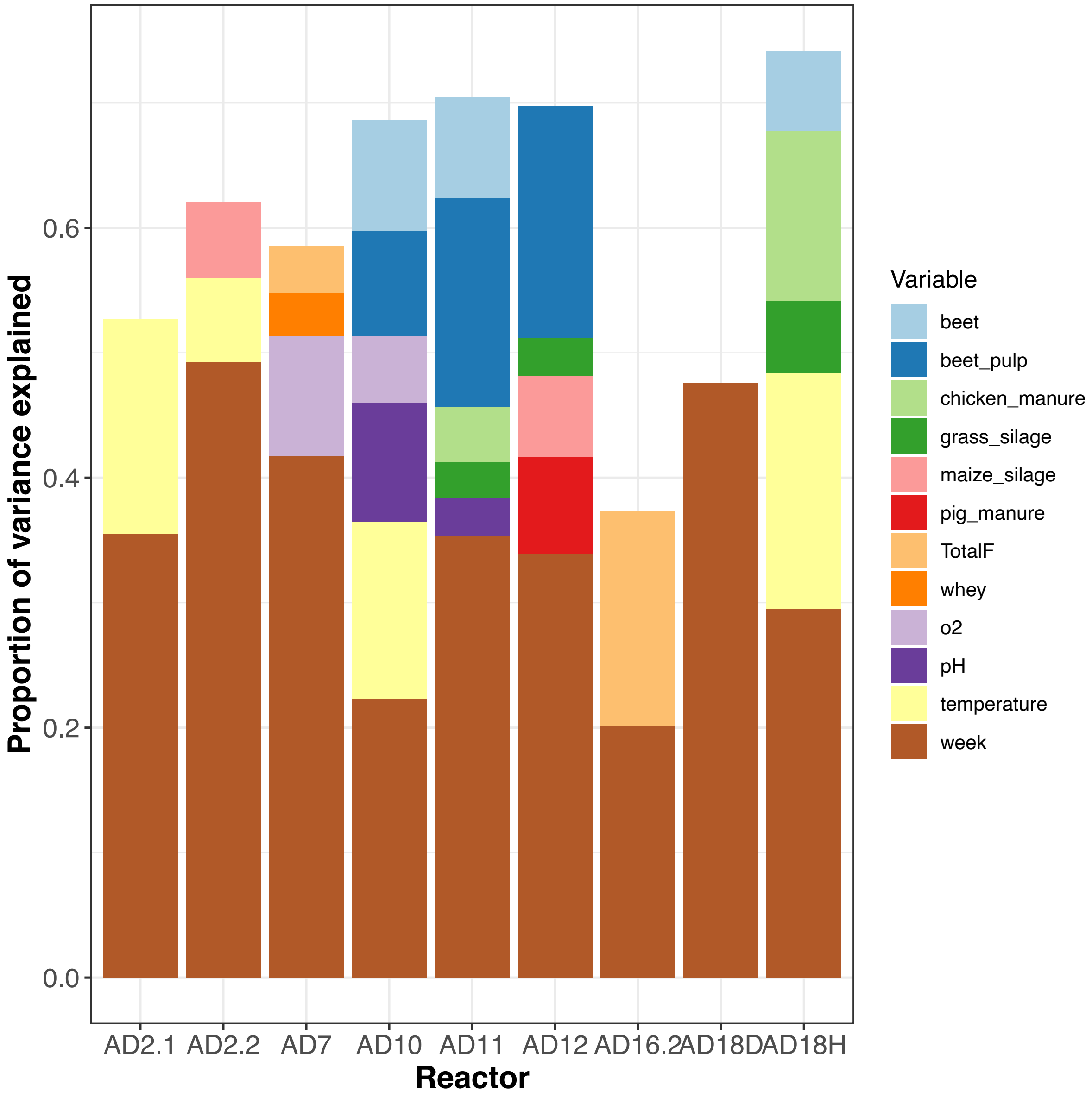


**Supplementary Figure 14. Proportion of variation in community explained by significant meta data variables for each reactor.** A multivariate permutation ANOVA (as implemented in the adonis function of vegan) of community structure (measured by normalised dMAG coverage) was performed against sampling week and all additional metadata variables that were at least 90% complete for that reactor. This was done for each reactor independently using Bray-Curtis distances. The bar chart shows the variance explained (R2) for those variables that were significant with $p < 0.05$. See also Supplementary Table 6.

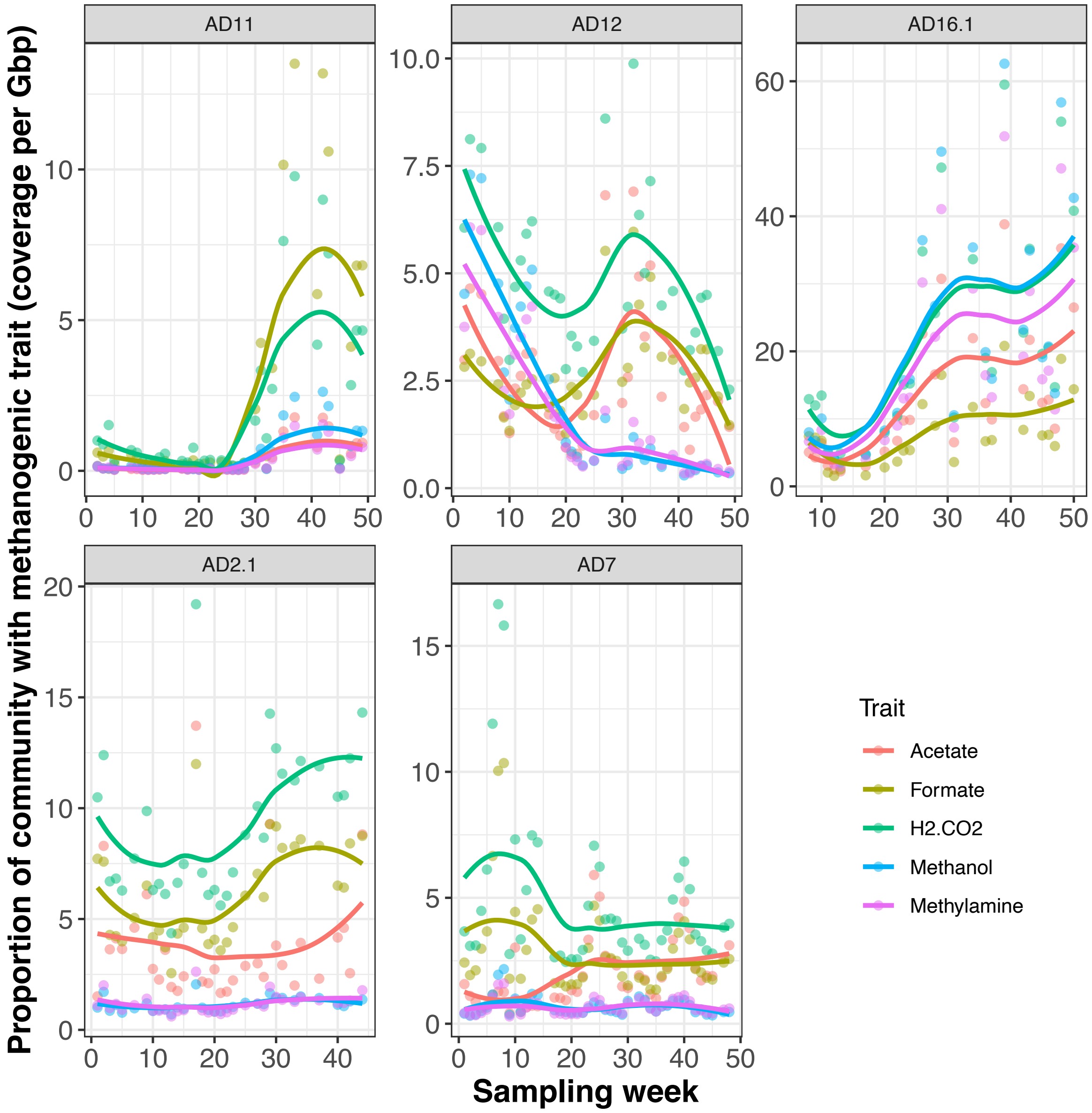


**Supplementary Figure 15. Changes in total predicted methanogenic substrate usage for each reactor over time.** To calculate methanogenic trait substrate usage abundance, we used the methanogenesis substrate assignments for each dMAG and summed the normalised abundance of dMAGs predicted to be capable of using a particular substrate (see Methods).

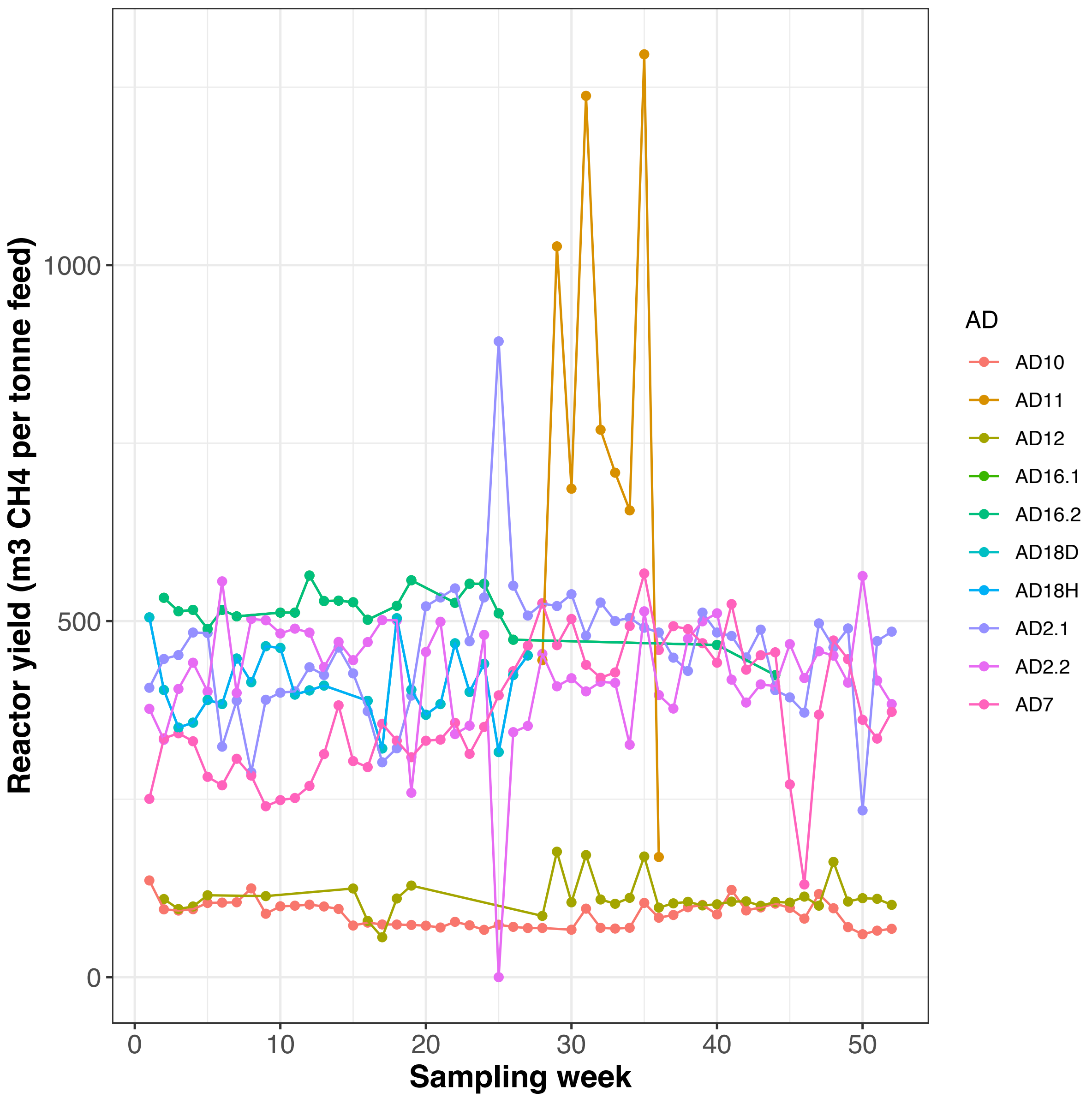


**Supplementary Figure 16. Changes in yield for each reactor over time.** Yield was defined as the total volume of methane gas (m3) produced per unit mass of feedstock (tonne).

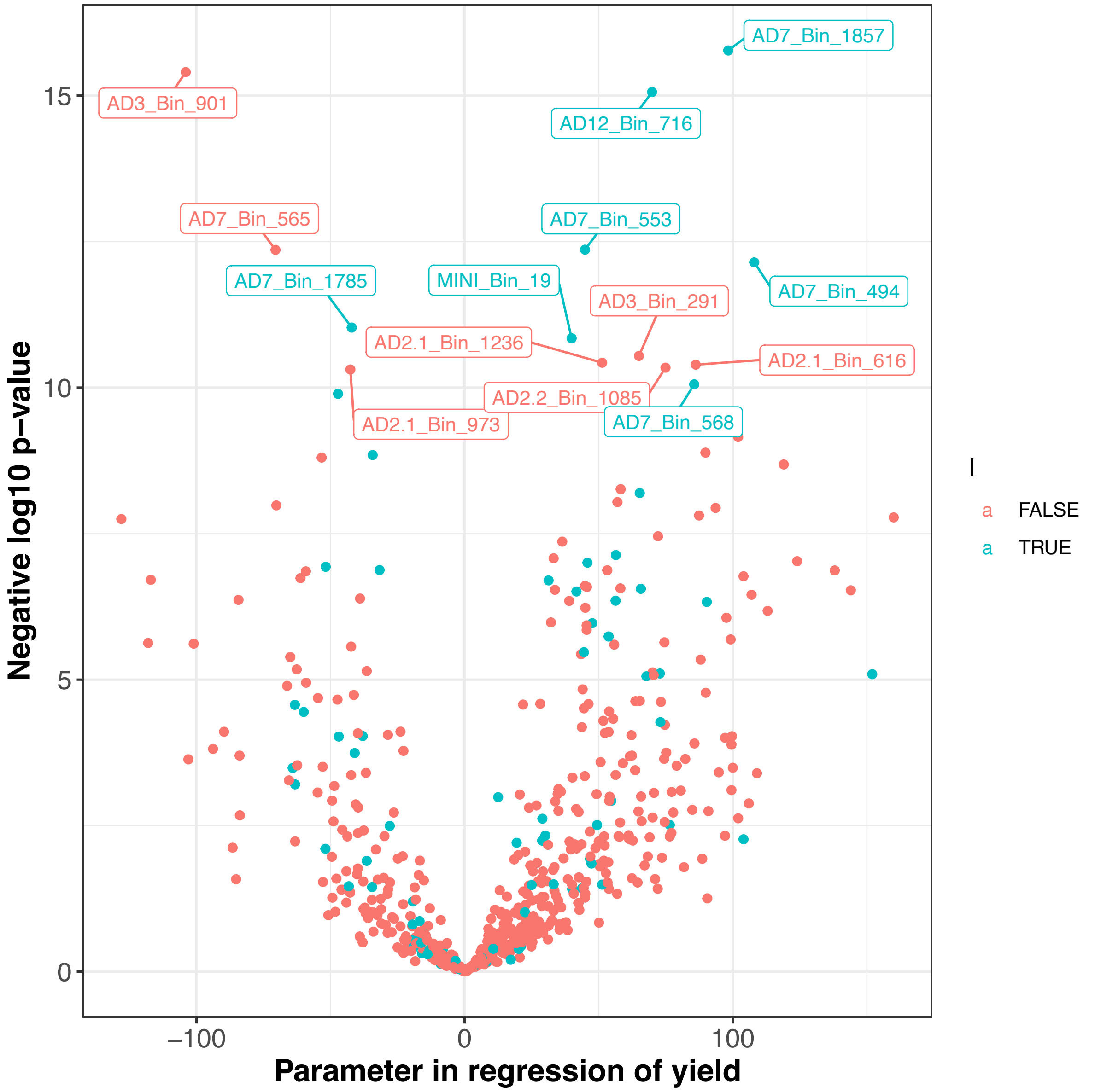


**Supplementary Figure 17. Volcano plot of yield against normalised dMAG coverages.** Each point on the plot shows a dMAG, the x-axis gives the multivariate regression coefficient of the log transformed dMAG coverage with yield accounting for reactor identity and operating temperature, the y-axis the negative log10 of the significance. Colour denotes whether a dMAG was predicted to be a methanogenic partner or not. See also Supplementary Table 9.
